# Uncovering the class II-bacteriocin predatiome in salivarius streptococci

**DOI:** 10.1101/2024.03.04.583286

**Authors:** Julien Damoczi, Adrien Knoops, Marie-Sophie Martou, Félix Jamaux, Philippe Gabant, Jacques Mahillon, Johann Mignolet, Pascal Hols

**Affiliations:** Biochemistry and Genetics of Microorganisms, Louvain Institute of Biomolecular Science and Technology, Université catholique de Louvain, Louvain-la-Neuve, Belgium; Syngulon, Seraing, Belgium; Laboratory of Food and Environmental Microbiology, Earth and Life Institute, Université catholique de Louvain, Louvain-la-Neuve, Belgium; Department of Fundamental Microbiology, Faculty of Biology and Medicine, University of Lausanne, Lausanne, Switzerland

**Keywords:** quorum sensing, predation, antimicrobial peptide, bacteriocins, ComRS, BlpRH, class II bacteriocin

## Abstract

Facing the surge of antibiotic resistance, the medical field has a critical need for alternatives to treat bacterial infections. Among these, the use of bacteriocins, ribosomally-synthesized antimicrobial peptides produced by bacteria, is considered to be a promising route. In the human commensal *Streptococcus salivarius*, the production of unmodified class II bacteriocins is directly controlled at the transcriptional level by the quorum-sensing ComRS system. Here, we used an integrated approach combining bioinformatics and synthetic biology to identify novel bacteriocins from salivarius streptococci active against human pathogens. We developed a bioinformatic pipeline that combines conservation of DNA motifs for genetic regulation and features of bacteriocin primary sequences to uncover cryptic class II bacteriocins. Notably, we discovered more than 50 novel bacteriocin candidates clustered into 21 groups from 100 genomes of *S. salivarius*. Strain-based analysis of bacteriocin cocktails revealed an important diversity restricted to seven distinct loci, probably resulting from bacteriocin intra- and inter- species exchanges. Using *in vitro* or *in vivo* production and synthetic biology tools, we showed that most of them are active against a panel of Gram-positive bacteria, including clinically-relevant pathogens. Overall, this work provides a new search-to-test generic pipeline for the discovery of novel bacteriocins in Gram-positive bacteria that could be used in cocktails for broad applications in the food and biomedical fields.

**IMPORTANCE:** To survive in highly challenging environments, streptococci have evolved a competence- predation coupling mechanism for genome plasticity. This developmental process is highly regulated at the transcriptional level, masking the predation killing effects in usual laboratory conditions. Here, we present a general strategy that combines bioinformatics and synthetic biology to unveil class II bacteriocins in streptococci. Its implementation to the beneficial species *Streptococcus salivarius* revealed around 40 class II salivaricin cocktails explained by the plasticity of seven independent loci. Notably, the salivaricin predatiome includes a subtle blend of fratricins, sobrinicins, and broad-spectrum bacteriocins with overlapping activities against a wide spectrum of low-GC Gram-positive bacteria, including notorious pathogens. Furthermore, most of those bacteriocins are predicted to be variants of a common α-hairpin structure, indicating that their mode of action evolved convergently. Finally, the discovery of *ca.* 50 novel bacteriocins offers perspectives for the rational assembly of potent cocktails active against pathogenic staphylococci, streptococci or enterococci.

## INTRODUCTION

Antimicrobial resistance is one of the biggest current challenges facing humankind. In 2019, an estimated 1,270,000 people died from drug-resistance infections (41). According to the World Health Organization, antibiotic resistance is one of the largest threats to global health. By 2050, more than 10 million deaths are forecasted due to antimicrobial resistance (43). With the surge of antibiotic resistance, there is a critical need for alternatives to treat bacterial infections. Diverse promising treatments are in development to curtail that phenomenon, such as bacteriophage therapy (9), probiotics (12), and RNA therapeutics (32). Another alternative is the use of bacteriocins (11, 29, 47). These small antimicrobial peptides, ribosomally synthesized by bacteria, are active against Gram-positive or Gram-negative bacteria but do not affect the producer, which typically encodes a specific immunity mechanism (16). So far, around a thousand bacteriocins are inventoried in the main bacteriocin database BAGEL4 (51). Yet, 99% of bacteria are predicted to produce at least one type of bacteriocin, implying that the bacteriocin reservoir available for bacterial infection treatment is massive (31).

Bacteriocins from Gram-positive bacteria are classified into two main groups: class I gathers peptides that undergo a range of post-translational modifications, while class II includes peptides that are unmodified or contain only minor modifications (*e.g.*, disulfide bridges) (1). Class II bacteriocins are subdivided in four subclasses: classes IIa, IIb, IIc, and IId that correspond to pediocin-like, two-peptide, leaderless, and non-pediocin-like single peptide bacteriocins, respectively (1). Except class IIc bacteriocins, class II bacteriocins are characterized by the presence of a leader sequence with a conserved addressing GG motif ([M|L|V]X_4_GG) (54). This sequence is essential for the secretion of the bacteriocin outside the cell and maintains the peptide in an inactive form to avoid intracellular toxicity issues (44). This work will focus on bacteriocins produced by *Streptococcus salivarius,* a Gram-positive lactic acid bacterium and a dominant commensal of the oral cavity (14). Several studies identified bacteriocinogenic strains of this species as highly potent against pathogenic bacteria. For that reason, several strains are already used as probiotics, such as M18 (7), K12 (15), and 24SMB (5). In *S. salivarius*, six class I bacteriocins (SlvA, SlvB, SlvD, SlvE, SlvG32, and Slv9) (2, 26) and six class II bacteriocins (BlpK, SlvV, SlvW, SlvX, SlvY, and SlvZ) (26, 38) have been identified so far.

In the genus *Streptococcus*, bacteriocin production is controlled by a cell-to-cell communication strategy known as quorum sensing (19, 38, 48). The regulation of class I and class II bacteriocins differs and is species-dependent. Class I bacteriocins as such act as stimulating pheromones and induce their own production through a phosphorylation cascade involving a dedicated two-component system (28, 34). In contrast, the production of class II bacteriocins is regulated by unmodified peptide pheromones that do not exhibit direct toxic effects (25). In streptococci, two regulation systems (*i.e.*, ComRS and BlpRH) control the production of class II bacteriocins (25). In *S. salivarius*, the ComRS system synchronizes competence and predation through upregulation of *comX* and bacteriocin genes, respectively (38). The precursor ComS is produced, secreted, and matured in the pheromone XIP (*comX*- inducing peptide). At a threshold concentration, XIP is internalized and interacts with the cytoplasmic sensor ComR (18, 20, 22). Then, the complex ComR-XIP activates transcription of target genes through its binding to a well-conserved DNA sequence called ComR-box (22, 38). This predation-competence coupling allows the bacteria to ensure the presence of free DNA in the environment before entering the competence state. In other streptococcal species, the production of bacteriocins is generally under the control of the BlpRH system (13, 19, 48), which is absent or inactive in *S. salivarius* (38). The pheromone precursor BlpC is produced, processed at a double-glycine motif, and exported by the dedicated transporter BlpAB (19). When the mature pheromone BlpC* reaches a threshold concentration, it activates the two-component system BlpRH (19). Upon activation, the trans-membrane histidine kinase BlpH phosphorylates the cytoplasmic response regulator BlpR, which in turn upregulates genes of the *blp* locus, including bacteriocin and self-immunity genes (13, 19, 36, 45).

Generally, the discovery of new bacteriocins is easier for class I compared to class II. Since class I bacteriocins and modification enzymes are encoded in the same locus, *in silico* analyses are based on the identification of putative bacteriocin and processing enzyme genes in close genomic vicinity (8, 40). In contrast, class II bacteriocins are generally encoded by small isolated (and usually unannotated) genes, either scattered throughout the genome (*e.g.*, *S. salivarius*) (38) or clustered in a unique locus (*e.g.*, *S. pneumoniae*) (36). Consequently, new bioinformatic pipelines need to be developed for their discovery.

In this work, we set up a combined approach based on bioinformatics and synthetic biology to identify novel class II bacteriocins in streptococci. We coupled the *in silico* recognition of regulatory DNA motifs with specific properties of known bacteriocins to unveil novel candidates. By testing our pipeline on *S. salivarius* genomes, we discovered 21 bacteriocin groups with multiple variants, including 13 groups never reported before. Using synthetic biology tools, we showed that unearthed bacteriocins display various profiles of inhibition against a range of Gram-positive bacteria, including a series of clinically-relevant pathogenic species (*e.g.*, *Staphylococcus aureus* and *Enterococcus faecium*).

## RESULTS

### *In silico* identification of 13 novel groups of class II bacteriocins in *S. salivarius*

To disclose novel salivaricins from a selection of 100 genomes of *S. salivarius* (complete or low contig number genomes), we first designed a multiparametric bioinformatic pipeline (Appendix S1 and Fig. 1). Considering that class II bacteriocins are hard to identify due to their small size and high sequence diversity, we combined three features: regulatory elements, sequence size, and signal sequence property. In several *S. salivarius* strains, the bacteriocin loci are under the direct control of the ComRS system (38). We therefore scouted the genomes for all open reading frames (ORF) that include a genuine ComR-box in their upstream genomic region (step 1) (Table S1). The ComR-box is a palindromic sequence oriented at the 3’ end by a T-rich stretch (T-track) (18, 20, 38). Therefore, we restricted the search to all ORFs downstream of the ComR-box T-track at a maximal distance of 4 kb (step 2). Then, we narrowed down the analysis by selecting sequences coding for precursor peptides ranging from 40 to 100 amino acids (step 3). We next discarded all the peptide sequences that do not include the conserved sequence motif ([M|L|V]X_4_GG) of class II bacteriocins (step 4). We finally retained the peptide sequences with specific lengths for leader sequences (from 14 to 28 amino acids) and associated mature sequences (from 25 to 70 amino acids) (step 5). Then, we manually removed all sequences devoid of a ribosome binding site upstream of the start codon (step 6) (Appendix S2).

**Figure 1.**
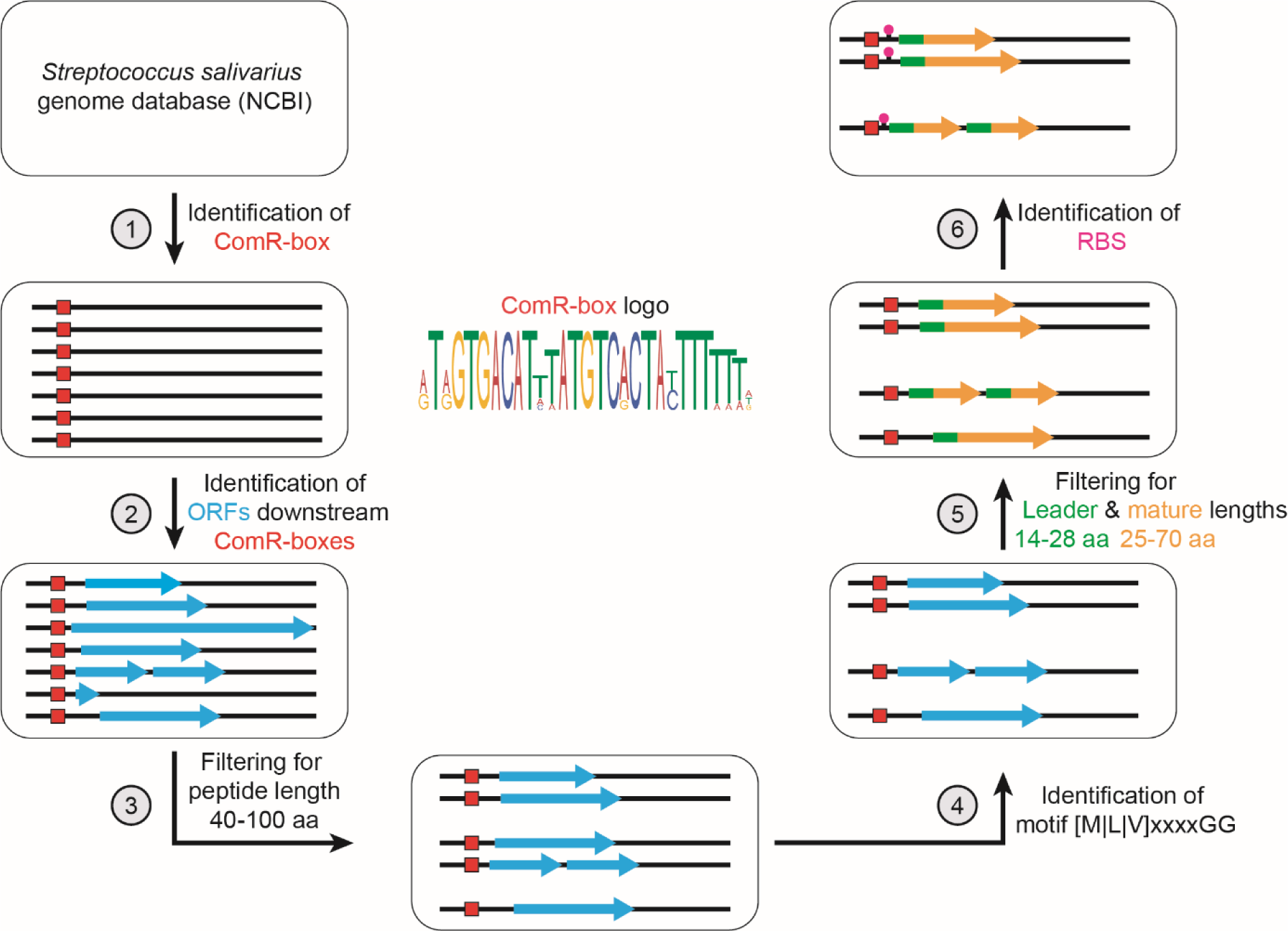
Schematic representation of the six steps of the multiparametric bioinformatics pipeline. Red boxes represent the ComR-box and blue arrows indicate the presence of downstream open reading frame(s). Leader sequences, mature sequences, and ribosome binding sites are shown as green squares, orange arrows, and purple dots, respectively.

We ultimately identified 21 putative bacteriocins, including six from *S. salivarius* (BlpK, SlvV, SlvW, SlvX, SlvY, and SlvZ; positive controls) and two from *Streptococcus thermophilus* (BlpE, and BlpF) that were previously reported (21, 38). Since *S. thermophilus* and *S. salivarius* are two closely related species, the identification of known bacteriocins from *S. thermophilus* suggests that they were exchanged through horizontal gene transfer. From this *in silico* analysis, we unearthed 13 novel bacteriocin candidates that were named PsnA to PsnM (<u>P</u>an <u>S</u>alivarici<u>n</u> A to M) (Table 1). We also noticed that several bacteriocins are always encoded in tandem (PsnA-PsnB, PsnC-PsnD, PsnE-PsnF, PsnG-PsnH, and SlvY-SlvZ), suggesting that they could belong to the two-peptide class IIb bacteriocins (42). Together, these results show that our bioinformatic pipeline can identify a range of novel class II bacteriocins in *S. salivarius*.

**Table 1.**
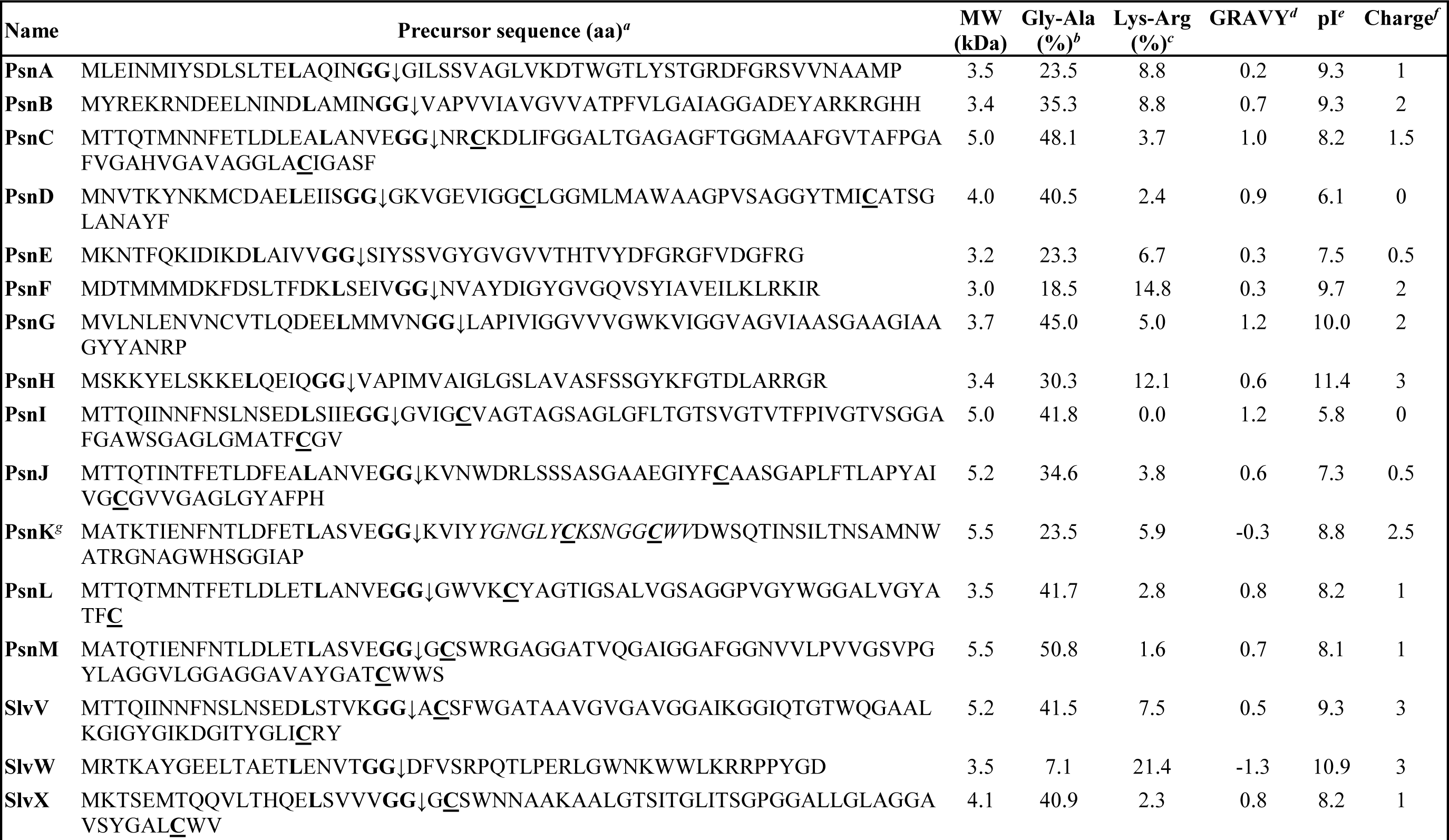
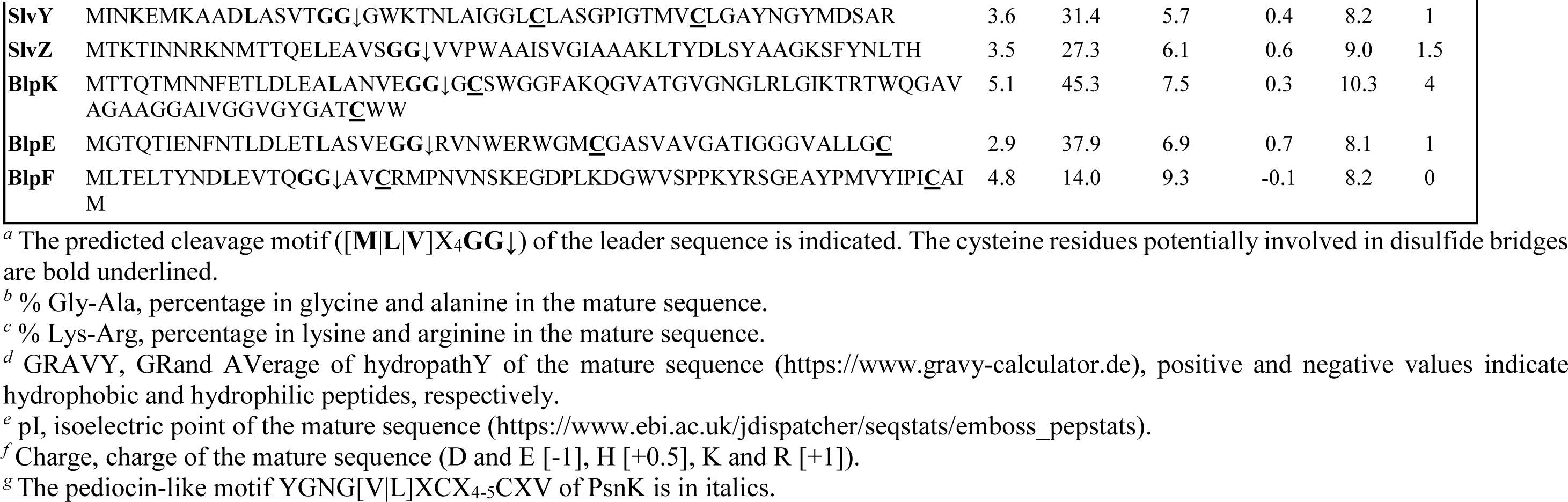
Class II bacteriocins of *S. salivarius* identified by *in silico* analysis.

### Class II salivaricins are variably distributed among streptococcal species

We next investigated the diversity of identified salivaricins across the genomes of *S. salivarius* and other streptococci that could be found in the oral cavity, such as *S. thermophilus*, *Streptococcus pyogenes*, and *S. pneumoniae* (Fig. 2A). We clustered bacteriocin candidates using BLAST searches and defined variants as homologous peptides with an identity ≥ 50% and at least one different amino acid. In total, we identified more than 50 salivaricin variants (Table S2). We observed that ∼50% of salivaricin peptides (or their variants) (11/21) are restricted to the species *S. salivarius,* while others are moderately to highly spread (*i.e.*, BlpK) among the three other streptococcal species (Fig. 2A). Strikingly, some bacteriocins are more represented in other species than in *S. salivarius*. For example, BlpK is present in ∼25% of *S. salivarius* strains, while it is found in ∼75% of strains from *S. thermophilus* and *S. pneumoniae* (Fig. 2A). Evolutionary mechanisms explaining specific bacteriocin selection for each species remain unclear but is likely to relate to inter-species competition inside the ecological niche. We also observed a high sequence conservation between variants of each salivaricin within the same species (Table S2). For example, BlpK variants display a negative correlation between the phylogenetic distance and the sequence conservation (Fig. 2B and C). This suggests that bacteriocin gene exchanges through horizontal gene transfer between distant streptococcal species are uncommon events.

**Figure 2.**
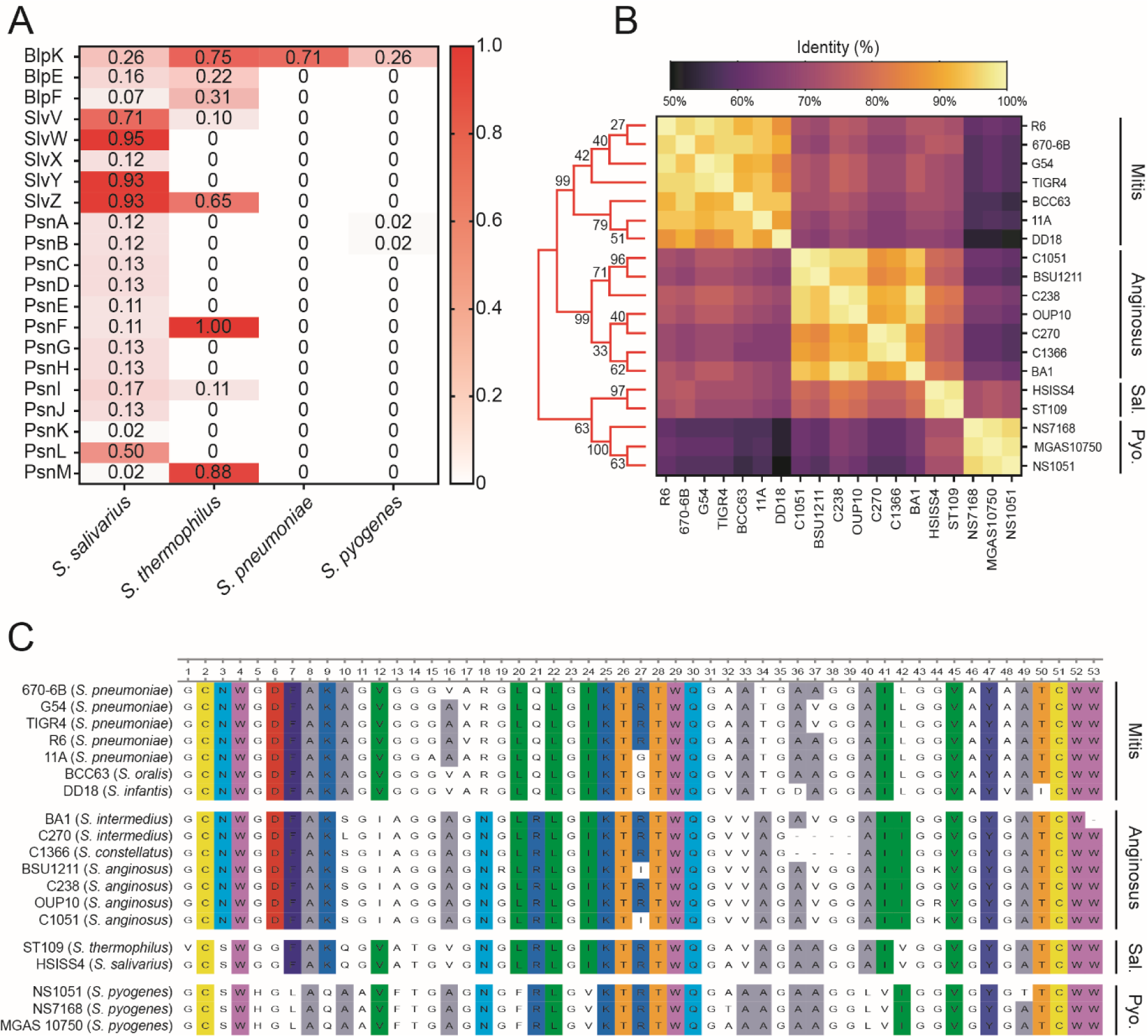
Distribution of salivaricins among oral streptococci. (A) Occurrence of the 21 prototypical salivaricins in *S. salivarius* compared to three streptococcal species that could be found in the oral cavity. The number in each box indicates the frequency of observation of the bacteriocin among all analyzed strains of the species. The analysis was performed on *S. salivarius*, *S. thermophilus*, *S. pneumoniae*, and *S. pyogenes* for 99, 78, 137, and 261 genomes, respectively. (B) Phylogenetic tree and heatmap of sequence identity (%) of natural BlpK variants from various streptococcal strains. The streptococcal groups are indicated on the right (Sal., salivarius; Pyo., pyogenic). BlpK sequences were aligned using MUSCLE and the tree was then generated with MEGA11 using the neighbor-joining method (10,000 bootstrap replicates). Bootstrap values are given at each node. (C) Alignment of natural BlpK variants. The alignment was generated with Clustal Omega and residues were colored (RasMol code) according to physicochemical properties.

Together, these analyses show that approximately half of class II salivaricins are restricted to *S. salivarius*, while the other half are variably shared through rare exchange events between streptococci that could be present in the oral cavity.

### Predatiome reshuffling results in a high diversity of salivaricin cocktails in *S. salivarius*

We also investigated the salivaricin gene content in all the *S. salivarius* strains that were used during the bioinformatic analysis (Fig. 3A and Table S3). First, we observed that 85% of all strains contain between five and seven bacteriocin genes (few strains can harbor up to 10 bacteriocin genes) arrayed in 37 different combinations (Fig. 3B). Second, we noticed that several specific bacteriocin genes are more widely distributed than others among *S. salivarius* strains. We identified that SlvY with SlvZ and SlvW are present in 100% and 95% of analyzed strains, respectively (Fig. 3C). In contrast, the bacteriocins PsnK and PsnM are only present in 2% of *S. salivarius* strains (Fig. 3C). We also observed a high co-occurrence for some bacteriocins (*e.g.*, SlvY with SlvZ, PsnA with PsnB), which supports their belonging to the two-peptide class IIb bacteriocins (Fig. 3A). Unexpectedly, we also observed bacteriocin gene exclusion (*e.g*., PsnL vs. BlpK, PsnI vs. SlvV) (Fig. 3A). This high intra-species variability in salivaricins prompts us to analyze in detail the genomic context of all bacteriocin loci (Fig. 4). We identified seven loci encoding various bacteriocins and immunity proteins, most of them being highly divergent among *S. salivarius* strains (Fig. 4A-G). Analyzing the most variable locus (Fig. 4A), we noticed that the genes *blpK*, *psnL*, *psnJ,* and *psnC-D* are located at the same position, explaining their mutual exclusion at a single genome level (Fig. 4A). Interestingly, we also spotted gene fragments encoding bacteriocin leader sequences without their mature part (Fig. 4A, orange arrows), probably resulting from intra-chromosomal deletion events between directly repeated bacteriocin genes. Bacteriocin loci are highly variable islands framed by two strictly conserved regions. Therefore, we hypothesize that surrounding conserved regions in combination with conserved leader-encoding sequences are involved in double recombination events, potentially mediated by natural transformation. This specific mechanism would account for salivaricin genes intra- (and possibly inter-) species exchanges.

**Figure 3.**
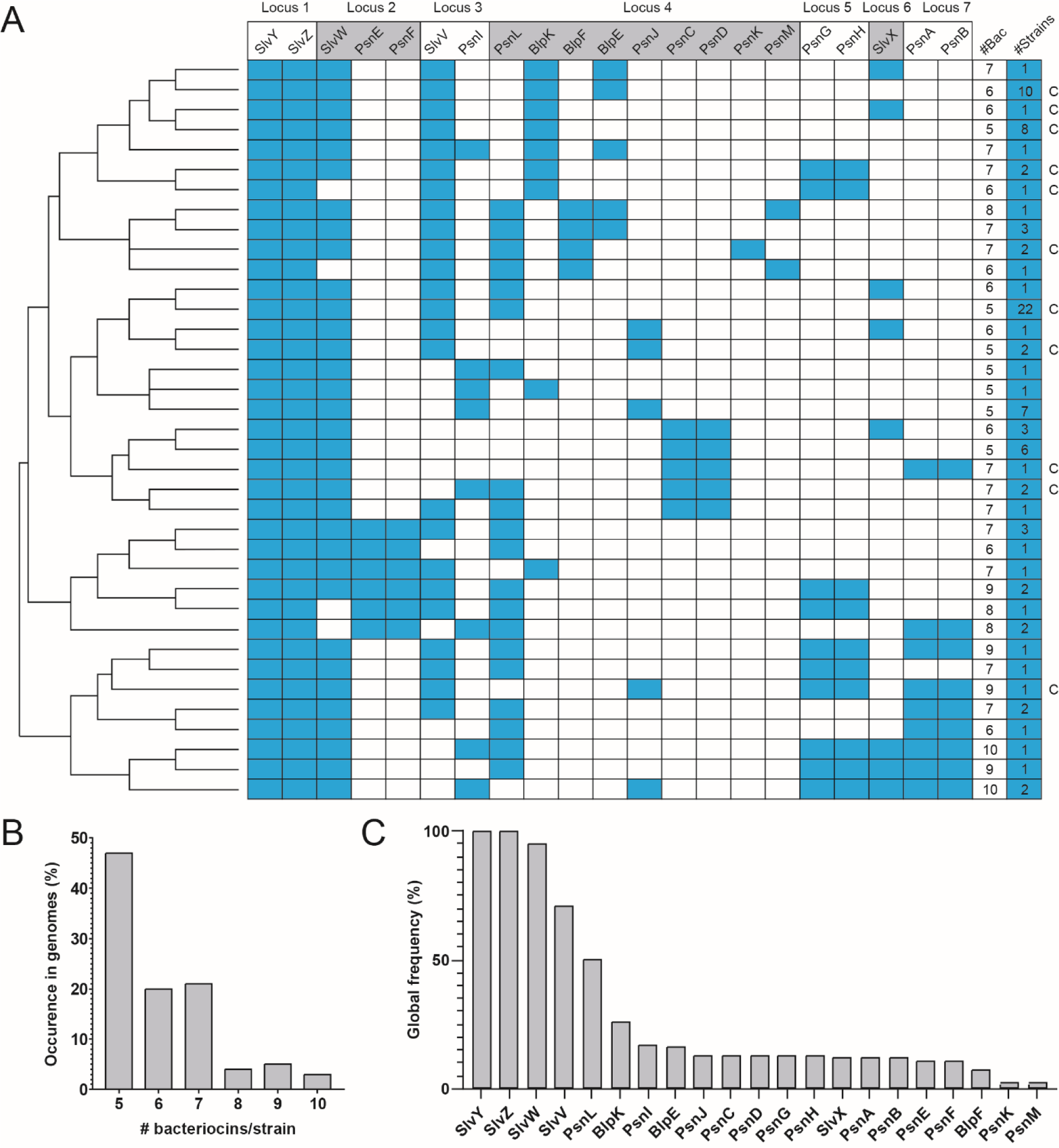
Composition of salivaricin cocktails identified in *S. salivarius*. (A) Schematic representation of the 37 salivaricin cocktails identified in seven loci from 100 *S. salivarius* genomes. A blue box indicates the presence of the bacteriocin. The hierarchical clustering in bacteriocin content of the different strains is shown on the left. The C letter on the right indicates that the cocktail was identified in at least one strain for which a closed genome is available. (B) Occurrence (%) of the number of salivaricins among *S. salivarius* strains. (C) Occurrence (%) of each individual salivaricin in *S. salivarius*.

**Figure 4.**
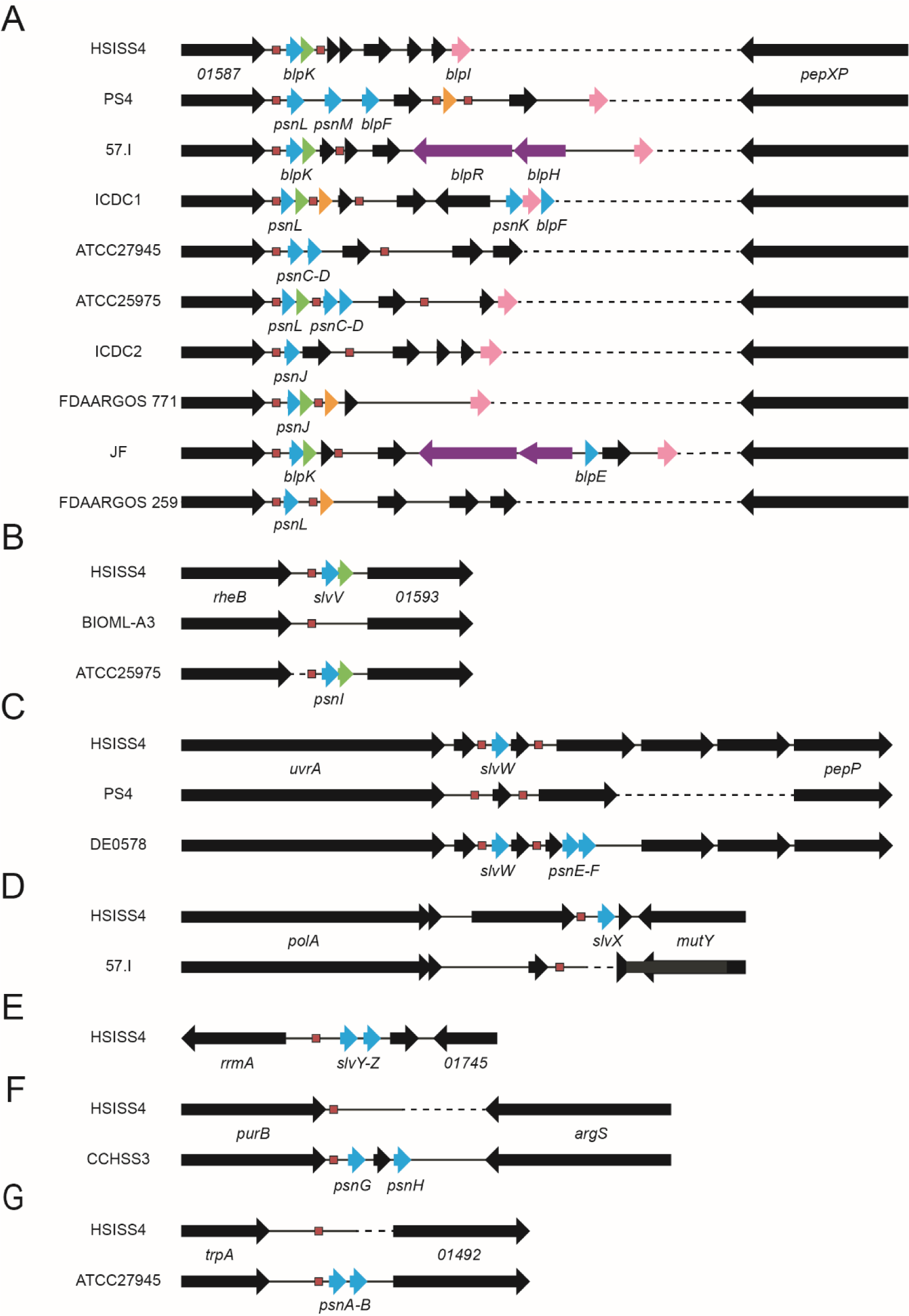
Comparison of the seven salivaricin loci in *S. salivarius.* Panels A to G display a representative variability in the seven different bacteriocin loci. A red box indicates the presence of a ComR-box; arrows represent genes encoding bacteriocins (blue), immunity peptides (green), bacteriocin leader sequences without the mature part (orange), fragments of the BlpRH system (purple), homologues of BlpI (pink), and other unrelated products (black). Dotted lines are spaces that were artificially added to align conserved genes in order to better visualize the genetic reorganization of each locus.

Together, these analyses demonstrate that the cocktail of class II salivaricins is highly variable at the species level and that this diversity partially relies on a specific locus, which is a potential hot spot of recombination.

### A majority of class II salivaricin candidates display antibacterial activity

Since our bioinformatics analysis identified 13 novel salivaricin candidates, we decided to evaluate their activity and prey spectrum. In a first step, we produce them in their mature form using synthetic DNA blocks coupled to an *in vitro* transcription-translation system, as reported before (24). Each cell-free sample was directly tested against three reporter strains: *Lactococcus lactis* IL1403, *Streptococcus thermophilus* LMD-9, and *S. salivarius* HSISS4. Antibacterial activity was identified for 5 out of 13 novel bacteriocin candidates (Fig. S1). In a second step, we decided to assess bacteriocin activity *in vivo* for all the 21 putative salivaricins. We chose to produce them directly in *S. salivarius* since class II bacteriocins may undergo minor post-translational modifications that require enzymes present in the original host. For instance, 60% of the salivaricin candidates display two cysteine residues, which might be involved in the formation of a disulfide bridge required for their functionality (Table 1) (21). To evaluate the activity of each individual bacteriocin in a single genetic background, we designed fusions between expression-secretion signals of the *blpK* gene and the mature part of each bacteriocin candidate (Fig. 5A). This strategy ensures tight control of expression through the XIP-inducible *blpK* promoter and secretion compatibility between the BlpK leader sequence and the bacteriocin exporter ComA (also XIP-inducible) in *S. salivarius* HSISS4 (Fig. 5A) (38). Those fusions were inserted at a permissive ectopic locus in a mutant strain devoid of any residual antibacterial activity (HSISS4 Δ*slv5*) (38).

**Figure 5.**
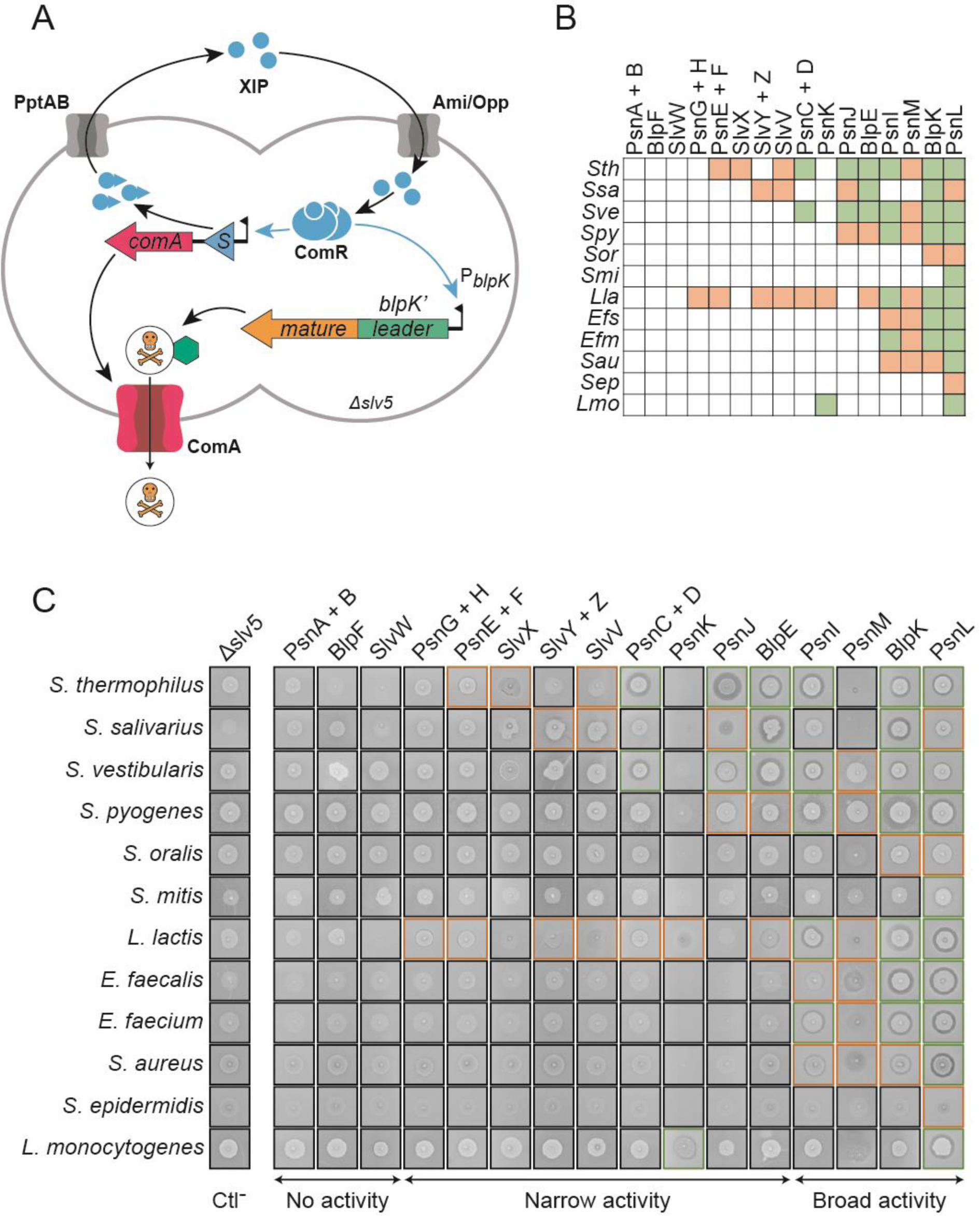
Activity assays performed with individual bacteriocins produced by *S. salivarius*. (A) Test strain (Δ*slv5*) for *in vivo* production of salvaricins. The gene fragment encoding the mature part of a salivaricin (orange arrow box) was fused to expression-secretion signals of *blpK* (P*_blpK_*-*blpK*’; green rectangle). The addition of the XIP inducer (blue circles) activates ComR, which will in turn stimulate the production of ComS (positive feedback loop via PptAB-mediated ComS export and Opp/Ami-mediated XIP import) and ComA (bacteriocin exporter, red arrow box). Concomitantly, the ComR-XIP complex activates the production of the hybrid bacteriocin by binding to P*_blpK_*. (B) Spectra of salivaricins against 12 Gram-positive bacteria belonging to the *Streptococcus*, *Lactococcus*, *Enterococcus*, *Staphylococcus*, and *Listeria* genera (list of strains in Table S5). Green, orange, and white boxes refer to high, weak, and absence of activity. (C) Spot-on-lawn assays with the 21 producer strains used to generate the table shown in panel B. Single-peptide producers of class IIa bacteriocins were mixed. High and weak activities are surrounded by green and orange lines, respectively. Producers are grouped into three categories: no activity, narrow spectrum, and large spectrum. Strain Δ*slv5* without bacteriocin activity is used as a negative control (Ctl^-^). Presented data were assembled from 40 independent plate assays. Each inhibition spot is representative of at least two independent experiments performed in the same conditions.

The collection containing 21 bacteriocin-producing strains was tested for antimicrobial activity against a broad range of Gram-positive bacterial species (including human pathogens) by spot- on-lawn assays. Since several bacteriocins belong to the two-peptide class IIb bacteriocins (PsnAB, PsnCD, PsnEF, PsnGH, and SlvYZ), we evaluated their activities by spotting a mixture of the two clones encoding each individual peptide (Fig. 5B). As control, we proved that these peptides have no activity when tested individually (Fig. S2). We observed that PsnI, PsnL, PsnM, and BlpK have the broadest spectrum of prey species. Those bacteriocins are active against closely related species but also against a group of clinically-relevant pathogenic bacteria more phylogenetically distant, such as *Staphylococcus aureus*, *Enterococcus faecium*, and *Listeria monocytogenes*. In contrast, several bacteriocins showcases a narrow spectrum of activity. The antimicrobial activity of PsnCD, PsnEF, SlvYZ, PsnJ, and BlpE targets only streptococcal species, while PsnK is mainly active against *L. monocytogenes.* The anti-listeria activity of PsnK correlates with the presence of a pediocin-like motif (YGNG[V|L]XCX_4- 5_CXV) in its mature sequence, which is a typical feature of class IIa bacteriocins active against the *Listeria* genus (Table 1) (17). Still, PsnAB, BlpF, and SlvW did not show any activity. We can hypothesize that they are inactive, non-bacteriocin peptides, not properly processed, or simply active against bacterial species outside of the tested spectrum.

Altogether, these results validate a bacterial chassis designed for testing class II bacteriocins *in vivo* and demonstrate antibacterial activity from a narrow to large spectrum for 17 out of 21 *in silico*-identified bacteriocin candidates.

### Enriched salivaricin cocktails potentiate their antibacterial activity

To determine how the composition of bacteriocin cocktails might affect the antibacterial activity of *S. salivarius*, we added PsnJ, PsnK, or PsnL to the native cocktail (BlpK, SlvV, SlvW, SlvX, and SlvYZ) of the strain HSISS4. While PsnL was chosen for its broadest spectrum of prey species, PsnJ and PsnK were selected for their more restricted activity against salivarius streptococci and *L. monocytogenes*, respectively (Fig. 5B). To generate the expanded cocktails, we transferred each of the three cognate fusions described in the previous section in the wild-type HSISS4 background. The three resulting strains were tested for their antimicrobial activity against the same panel of 12 prey species by spot-on-lawn assays (Fig. 6 and Fig. S3). Although the inhibitory activity of the expanded cocktails remains unchanged against most species compared to the native cocktail produced by HSISS4 (Fig. S3), we observed that the addition of a single bacteriocin could increase its activity against specific species (Fig. 6). First, the addition of the narrow-spectrum bacteriocin PsnK or PsnJ strongly increased activity against *L. monocytogenes* or *S. thermophilus*, respectively (Fig. 6). Notably, the anti-listeria effect of PsnK is clearly synergistic with the native cocktail, while the impact of PsnJ seems to be more limited (Fig. 6). As PsnK is the only member of class IIa among all identified salivaricins, its specific mode of action could explain this synergy with at least one native bacteriocin, which have no or weak anti-listeria activity when produced alone (Fig. 5B). Second, the addition of the broad-spectrum bacteriocin PsnL increased the cocktail efficiency against the human pathogens *S. aureus* and *L. monocytogenes*, and the human opportunist *Staphylococcus epidermidis*. In contrast to PsnK that synergize with the native cocktail of HSISS4 strain, PsnL alone accounts for most of the anti-staphylococci and anti-listeria activity (Fig. 5B).

**Figure 6.**
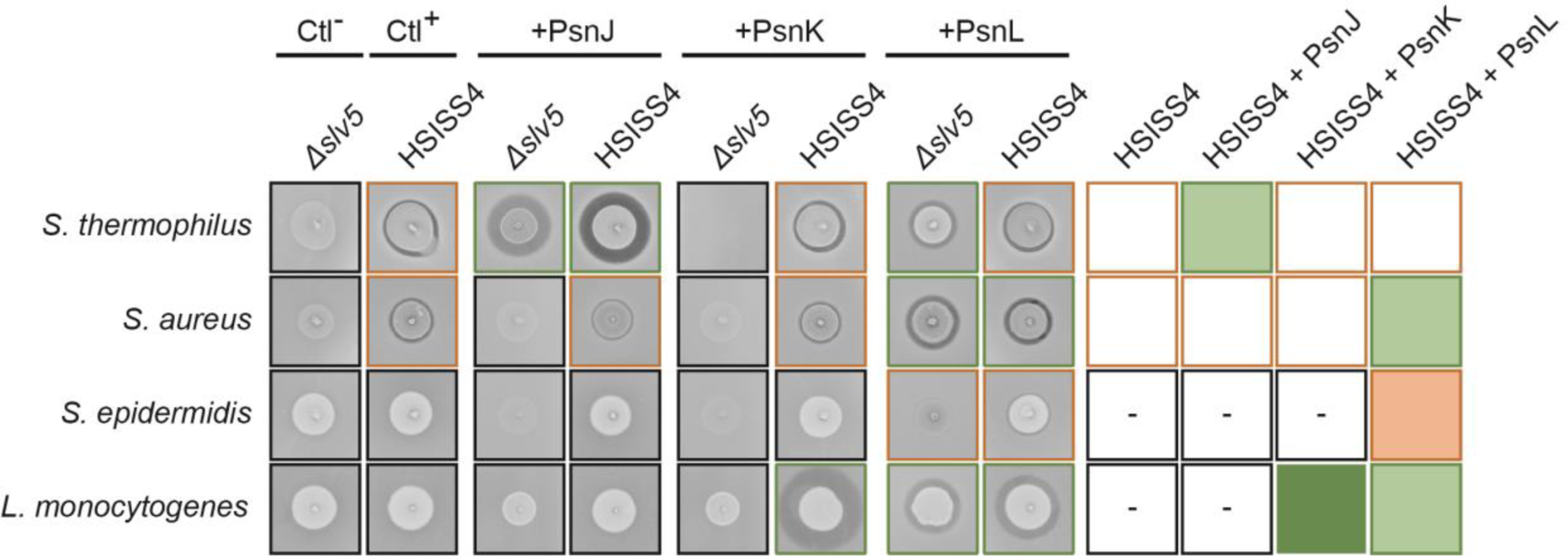
Activity assays of modified salivaricin cocktails produced by *S. salivarius* HSISS4. Spot-on-lawn assays (left panels) of HSISS4 derivatives (wild-type and Δ*slv5*) producing PsnJ, PsnK, or PsnL against four Gram-positive bacteria (list of strains in Table S5). Strains Δ*slv5* and HSISS4 (wild-type) are used as negative (Ctl^-^) and positive (Ctl^+^) controls, respectively. High and weak activities are surrounded by green and orange lines, respectively. A schematic representation of inhibitory spectra from HSISS4 cocktails is shown in the right panel. Boxes with a minus sign indicate that no activity was detected. Boxes surrounded in green and orange indicate high and weak activity, respectively. Boxes filled in green and orange indicate a modified activity due to the presence of the additional salivaricin in the cocktail of wild-type HSISS4; light and dark colors refer to additive and synergic effects, respectively. Each inhibition spot is representative of at least two independent experiments performed in the same conditions.

Together, these results underline the importance of salivaricin combinations for enhanced efficiency and extended anti-bacterial spectra, as illustrated here for growth inhibition of clinically-relevant pathogens.

## DISCUSSION

In the past, novel bacteriocins were discovered by large-scale screening of producer candidates in combination with prey species of interest using the cell growth defect as a readout, followed by bacteriocin purification and identification of their sequences by mass spectrometry. This approach is not only time-consuming but also very restrictive for bacteriocins that are produced in laboratory conditions. This is particularly true for Gram-positive bacteria, and specifically streptococci, where the production of most bacteriocins is strictly induced by a range of transcriptional regulatory systems (RNPP cytoplasmic sensors and/or TCSs) (25, 26, 48). Hence, the whole antimicrobial potential of a defined species remains mostly underestimated. To unveil this potential, various bioinformatics approaches have been undertaken, but none of them include the specific feature of genetic regulation as a key parameter to identify functional bacteriocin genes. The addition of this parameter to specific bacteriocin features was expected to strongly increase the power and sensitivity of the bioinformatic analysis (lower number of false negatives and positives). In this work, we fully validated this approach by unveiling the *S. salivarius* predatiome of active class II bacteriocins under the control of the ComRS system. We also benchmarked our approach with classical bioinformatic tools used to identify bacteriocins in bacterial genomes, such as BAGEL4 (51) and AntiSMASH (6). Because those tools are searching for the synteny between bacteriocin genes and genes encoding post-modification enzymes, immunity factors, or export systems, they failed to identify ∼ 50% of the class II bacteriocin genes in *S. salivarius* genomes. Since the identification of bacteriocin candidates in *S. salivarius* was successful with our bioinformatic pipeline, we quickly evaluated if it could be applied to other streptococcal species whose bacteriocin genes could be controlled by the ComRS system (Table S1). We selected one representative genome of three species (*i.e.*, *Streptococcus vestibularis*, *Streptococcus downei*, and *Streptococcus sobrinus*) and identified multiple bacteriocin candidates in each genome (Table S4). While bacteriocin candidates of *S. vestibularis* were all previously identified in *S. salivarius*, 12 uncharacterized bacteriocins were discovered in the genomes of the two other species (Table S4). Additionally, we also implement our bioinformatic approach to identify streptococcal bacteriocins that could be regulated by the alternative signaling system BlpRH (Table S1). As reported above, we used one representative genome of three species (*i.e.*, *Streptococcus pneumoniae*, *Streptococcus oralis*, and *S. thermophilus*) whose bacteriocin genes are predicted to be controlled by the BlpRH system. From the 19 bacteriocin candidates identified in the three genomes, three bacteriocins from *S. oralis* were never reported before (Table S4). These two additional *in silico* analyses show that our multiparametric bioinformatic tool could be broadly applied for the discovery of novel class II bacteriocins in a range of streptococcal species. Finally, we reckon that this regulation-based screening could be adapted to other quorum sensing systems (or sensory modules) in many bacteria to drastically increase the identified reservoir of bacteriocins and associated functions (*e.g.*, maturation enzymes, immunity proteins, signaling peptides).

The *in silico* analysis of the *S. salivarius* predatiome of class II bacteriocins revealed interesting features in terms of inter- and intra-species distributions. At the inter-species level, the class II salivaricins are partially shared between the three streptococcal species of the salivarius group. For instance, ∼ 40% of salvaricins are common to *S. salivarius* and *S. thermophilus* with a high level of protein identity (Fig. 2A and Table S2). Since these two species cross-talk by using very similar XIP pheromones and display a high nucleotide identity at the genome level (90- 95%) (14, 18), the bi-directional spreading of bacteriocin genes could be strongly favored by natural transformation. By contrast, their low distribution in other streptococcal groups (Fig. 2A) is certainly constrained by the absence of cross-talk and lower genomic sequence conservation (49). At the intra-species level, we observed a high variability of the salivaricin content (Fig. 3). While class II bacteriocins were reported to be mainly grouped in a single cluster (*blp* locus) in *S. pneumoniae* and *S. thermophilus* (19, 36), they are scattered throughout seven unlinked loci in *S. salivarius* (Fig. 4). This large spread at the genome level favors a high variability of bacteriocin cocktails (∼ 40 over 100 strains). However, the *blp*-like locus (locus 4), which contains pseudogenes of the BlpRH system (38), remains the more dynamic locus for bacteriocin gene insertion, deletion, or swapping. Indeed, it could be considered as a plasticity platform for bacteriocin gene exchanges as it encodes nine different bacteriocin peptides and up to four in the same strain (Fig. 3A and 4).

Our systematic work of testing the activity of each individual bacteriocin peptide sheds light on its respective contribution to bacteriocin cocktails in different strains. In *S. salivarius*, competence and predation are intimately interlinked since both processes are directly controlled by the ComRS system (38). Competence-induced bacteriocins that kill bacteria of the same species or closely related species could be considered as fratricins or sobrinicins, respectively (10, 27, 33, 53). Both types of killing peptides (in association or not with bacteriolysins) are able to release exogenous DNA by lysis from neighboring bacteria. Next, free DNA could be captured by the transformation machinery with an opportunity of genome integration, depending on the DNA homology of the prey genetic material. The repertoire of class II salivaricins active against streptococci includes both fratricins and sobrinicins (Fig. 7A). On one hand, SlvYZ can be considered as a strict fratricin that only targets *S. salivarius* among streptococci. The targeting of siblings seems to be a basic function of the competence-predation coupling since this bacteriocin is present in all *S. salivarius* genomes analyzed during this study (Fig. 3A and 5B). On the other hand, SlvX, slvV, PsnCD, and PsnEF can be seen as narrow-range sobrinicins targeting species of the salivarius group, while PsnJ and BlpE are large-range sobrinicins with an extended repertoire against other streptococcal groups (Fig. 5B and 7A). Interestingly, *S. salivarius* is also equipped with four broad-spectrum bacteriocins of gradual increased activity (*i.e.*, PsnI < PsnM < BlpK < PsnL) against many genera among *Firmicutes* (low-GC Gram-positive bacteria) (Fig. 5B and 7A). Such a large spectrum of prey species may suggest that their roles overcome the competence-predation coupling for genome plasticity. The high occurrence (around 85%) of at least one of these large-spectrum bacteriocins in *S. salivarius* strains suggests an additional role in general predation for niche occupancy, which is in correlation with the high incidence of this species as a commensal in many sub-niches of the human body (*e.g.*, oral cavity, upper airways, and upper part of the small intestine) (30, 35, 50). As *S. salivarius* is present as a consortium of multiple strains (50), its shared predatiome of class II bacteriocins revealed a combinatory strategy of overlapping antimicrobial activities from a (very) narrow to large spectrum against *Firmicutes* (Fig. 7A). Besides a wider inhibition of competitors, the selective pressure that has shaped overlapping activities against a defined species is most probably dictated by an increased inhibitory efficacy (*i.e.*, additive and/or synergistic effects), as shown here by the mimetic transfer of novel bacteriocins in a wild-type background (Fig. 6).

**Figure 7.**
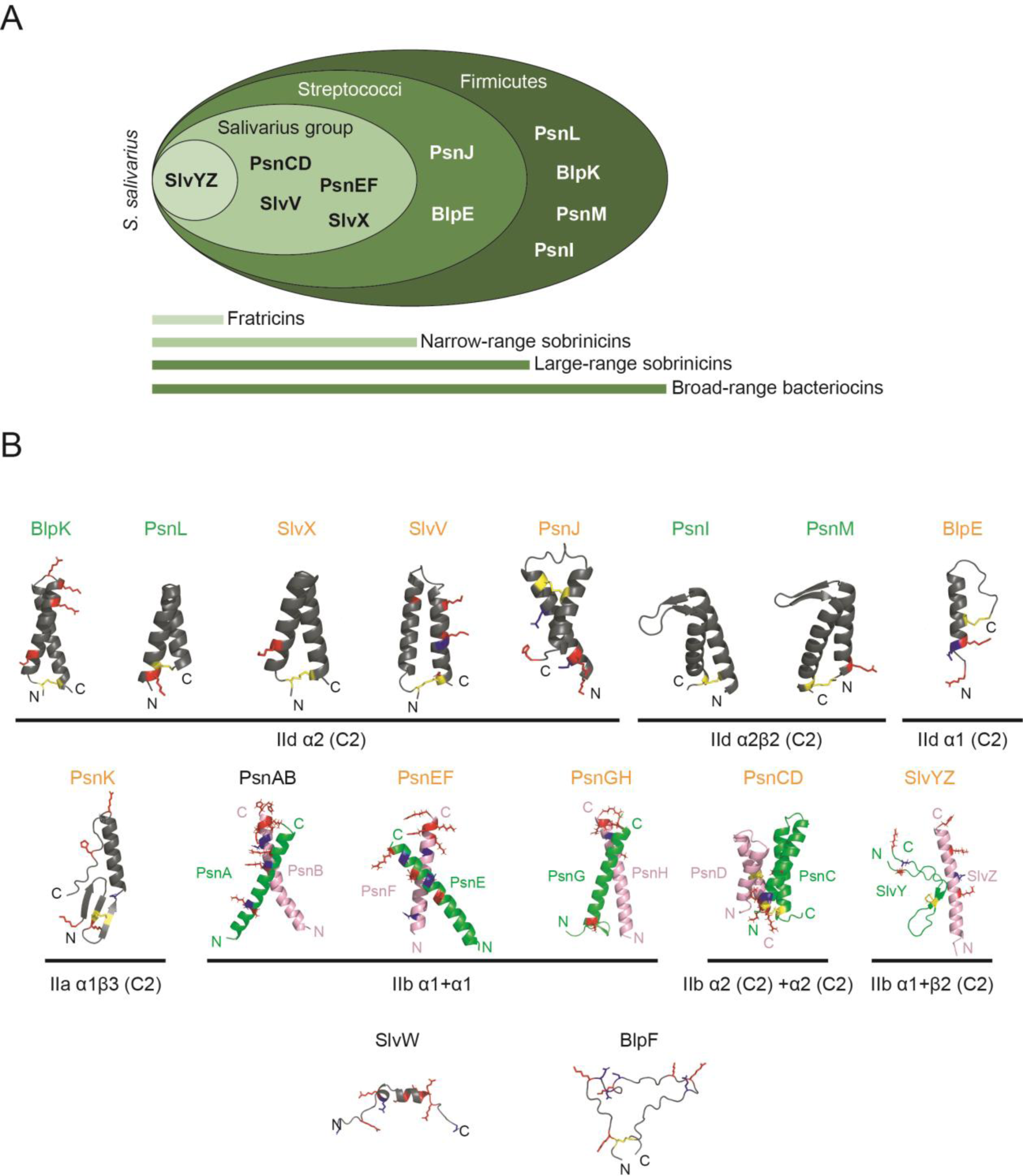
Global prey spectra and structure prediction of class II salivaricins. (A) Global overview of the overlapping activity spectra of active salivaricins against streptococci and *Firmicutes*. The activity against *L. lactis* (dairy strain) is not included here since considered as not relevant for the ecology of the digestive tract. (B) Structure prediction by AlphaFold 2 of monopeptides and two-peptide bacteriocins. Predicted 3D structures are organized according to their subclasses (IId, IIb, and IIa) in class II bacteriocins and shared structural elements (α helices and β strands). The presence of a predicted disulfide bond (C2) is also indicated. The color of the bacteriocin name in green, orange, and black refers to a large spectrum of prey species, a narrow spectrum of prey species, and the absence of identified activity, respectively. Monopeptides are in dark gray, while dipeptides are in green and pink. Cysteine, positively charged, and negatively charged residues are colored in yellow, red, and blue, respectively.

To better understand their evolution and mode of action, we also predicted the 3D structure of salivaricins using AlphaFold 2 (Fig. 7B) (37, 39, 52). Among class IId bacteriocins (single peptide), which include the largest set of active salivaricins, most are predicted to be α-hairpin peptides formed of two anti-parallel α-helices stabilized by a disulfide bond (3). Although initially considered class IId bacteriocins in our study, BlpF and SlvW are not or weakly structured in correlation with an absence of detected antimicrobial activity (Fig. 5 and 7). We hypothesized that they could be either helper antimicrobial peptides or uncharacterized signaling peptides. The predicted structures of most class IIb heterodimeric bacteriocins mimics α-hairpin peptides (*i.e.*, PsnAB, PsnEF, and PsnGH) (Fig. 7B) or are the association of two α- hairpin peptides (*i.e.*, PsnCD) as recently reported for the atypical class IIb Gallocin A (46). Finally, the predicted structure of PsnK is in full accordance with the well-established structures of pediocin-like bacteriocins of class IIa, shown to use the Man-PTS system as a receptor (23, 55). With some exceptions (*i.e.*, BlpE, SlvYZ, and PsnK), the structures of most class II salivaricins are variations or mimetics of α−hairpins, highlighting a possible convergent evolution for their activity. Another feature shared by the majority of salivaricins is the presence of positively-charged residues, which could be clustered in the α-hairpin linker region (*i.e.*, BlpK) or at C-terminal extremities (*i.e.*, PsnAB, PsnEF, and PsnGH) (Table 1 and Fig. 7B). As previously reported (4), we hypothesized that those residues are key to initiate interactions with negatively charged lipids in the plasma membrane. Except PsnK that is presumed to interact with the Man-PTS, the mode of action of class II salivaricins such as the requirement of a specific receptor or the ability to oligomerize for pore formation will require further investigations.

To conclude, we set up a novel combined approach for bacteriocin discovery that integrates their mode of regulation, exploited here for both *in silico* analysis and *in vivo* validation of their activities. This approach allowed us to extensively reveal the predatiome of class II bacteriocins in *S. salivarius* and offers a high potential to unveil the comprehensive predatiomes of many Gram-positive species that are so far extremely underestimated. This generic approach is key to enrich bacteriocin databases and physical collections of synthetic genes such as PARAGEN (24) and maximally exploit the whole potential of these antimicrobial peptides for many future biocontrol applications. The characterization of the *S. salivarius* predatiome as such also offers many opportunities of development in the probiotics field, such as the prediction of the antimicrobial potential of existing bacteriocin cocktails in various strains or the design of second-generation probiotics with adapted cocktails targeting specific pathogens. Class II bacteriocins as unmodified peptides are easy to synthesize chemically and to engineer by genetic modification for optimizing their efficiency and/or activity spectrum (26). Hence, the antimicrobial spectra of BlpK, PsnL, PsnI, PsnM, and PsnK against clinically-relevant pathogens and the identification of their numerous natural variants should have a high potential for future exploitation in the food industry and biomedicine.

## MATERIAL AND METHODS

### Bacterial strains and DNA material

Bacterial strains, plasmids, oligonucleotides, and PCR fragments used in this study are listed in Tables S5, S6, S7, and S8 of the Supplemental Information, respectively.

### Growth conditions

*Streptococcus salivarius* HSISS4 and derivates were grown without shaking in M17G (glucose 1% [w/v]) medium (Oxoid) or CDMG (chemically-defined medium with glucose 1% [w/v]) (35) at 37°C. *Lactococcus lactis* IL1403, S*treptococcus vestibularis* F0396, *Streptococcus thermophilus* LMD-9, *Streptococcus pyogenes* 4549, *Streptococcus oralis* Si0464, *Streptococcus mitis* LMG 14557, *Staphylococcus aureus* ATCC 6838, *Staphylococcus epidermidis* LMG 10273, *Enterococcus faecium* ATCC 19434, *Enterococcus faecium* Si0159, and *Listeria monocytogenes* ATCC 51777 were all grown without shaking in M17G at 37°C, except *L. lactis* IL1403 that was grown at 30°C. Solid plates inoculated with *S. salivarius* were incubated anaerobically (AnaeroGen 2.5L, Thermo Scientific) at 37°C. Spectinomycin was added when needed at 200 µg/ml. Synthetic peptide sXIP (purity of 95%) was supplied by Peptide2.0 Inc. (Chantilly, VA, USA), resuspended in water, and used at a concentration of 250 nM, except if otherwise stated.

### Strain construction by natural transformation

*S. salivarius* HSISS4 was transformed with Gibson assembly products composed of three PCR fragments: (i) the upstream region of *tRNA^Ser^* associated with a *blpK* gene fragment containing expression signals and encoding the leader sequence of BlpK; (ii) the downstream region of *tRNA^Ser^* associated with the *spc* cassette (spectinomycin resistance); and (iii) the mature sequence of a bacteriocin. PCR fragments were amplified with the Q5 polymerase (NEB, Ipswich, MA, USA), following the manufacturer’s recommended protocol. To induce natural transformation, an overnight CDMG preculture was diluted in 500 μl of fresh CDMG at a final OD_600_ of 0.05 and incubated for 135 min at 37°C. Then, 1 µM of sXIP and linear DNA (Gibson assembly product) were added to the culture, and cells were further incubated for 3 h at 37°C before plating on M17G agar supplemented with spectinomycin. After transformation, all constructions were verified by DNA sequencing.

### Spot-on-lawn assays for bacteriocin activity

To detect bacteriocin activity, 50 µl of overnight cultures of producer strains were diluted in 1 ml of M17G and grown for 3 hours at 37°C (final OD_600_ of 0.5). In parallel, a first feeding layer (30 ml) of M17G 1.5% agar supplemented with XIP (250 nM) was cast on a plate. Then, a second layer (15 ml) of M17G 0.3% agar containing the indicator strain was poured on top of it. This layer contains 800 µl of an overnight culture of the indicator strain. On the top of this last layer, 3 µl of bacteriocin producers were spotted. Finally, the plate was incubated overnight in anaerobic conditions at 37°C.

### *In vitro* production of bacteriocins

Synthetic plasmids (pUC57 derivative) expressing the mature parts of bacteriocins under the control of the T7 promoter were provided by GenScript (Piscataway, NJ). These plasmids (Table S6) in combination with the cell-free system PURExpress (NEB) were used to produce bacteriocins *in vitro*. To synthesize bacteriocins, the reaction mixture was assembled on ice in the following order: 4 µl of solution A, 3 µl of solution B, 2 µl of nuclease-free water, and 1 µl of template DNA (100 ng/µl). Then, the reaction mixture was incubated for 3 hours at 37°C, and the activity was measured by a spot-on-lawn assay.

### Multiparametric bioinformatics script

The script used to perform the bioinformatic analysis was written in R (www.r-project.org). The code of this script and the data set obtained from the screening of 100 genomes of *S. salivarius* are available in Appendices S1 and S2, respectively. For *S. salivarius*, a detection limit of ComR-boxes in the upper 75% of identity to the consensus of the identified ComR-box (Table S1) was set to avoid identifying pseudo-ComR-boxes. In addition to the script, tBLASTn was used on each strain of *S. salivarius* to confirm bacteriocin presence and identify variants. The tBLASTn searches were performed with the default setting, except for the expect threshold parameter, which was increased to 5 to avoid false negative hits.

### 3D structure prediction of bacteriocins

Structure predictions of bacteriocins with AlphaFold 2 were obtained from the AlphaFold CoLab notebook (39, 52).

## AUTHOR’S CONTRIBUTION

J.D., P.G., J.Mi. and P.H; conception and design. J.D., M.S and F.J.; acquisition of data. J.D. and A.K.; creation of the bioinformatic script. J.D., J.Mi. and P.H.; analysis and interpretation of data. J.D., P.G., J.Mi., J.Ma. and P.H., draft or revising the manuscript. All authors read and approved the final manuscript.

## ACKNOWLEDGMENTS

The work of PH was supported by the Belgian National Fund for Scientific Research (FNRS, grants PDR T.0110.18/T.0111.22 and CDR J.0090.21), the Concerted Research Actions (ARC, grants 17/22-084 and 22/27-120) from Federation Wallonia-Brussels, and the Syngulon company. JD and AK held doctoral fellowships from FNRS (FRIA fellowship). FJ was supported by SWP Research from Walloon Region (grant N° 8065 Staphcontrol). JMi received funding from the European Union’s Horizon 2020 research and innovation program (Marie Skłodowska-Curie grant N° 101018461). PH is Research Director at the FNRS.

## SUPPLEMENTAL INFORMATION

Supplemental information includes 8 tables, 3 figures, and 2 appendices.

