## Supplemental Table S1-S8 and Fig S1-S3 for "Uncovering the class II-bacteriocin predatiome in salivarius streptococci"

#### 1. SUPPLEMENTAL TABLES

**Table S1. ComR and BlpR binding sites used for *in silico* bacteriocin searches in streptococci**

**Table S2. Natural variants of class II salivaricins**

**Table S3. Distribution of class II bacteriocins in *S. salivarius* genomes**

**Table S4. Bacteriocin candidates identified by *in silico* analysis in other streptococci**

**Table S5. List of bacterial strains used in this study**

**Table S6. List of plasmids used in this study**

**Table S7. List of oligonucleotides used in this study**

**Table S8. List of PCR fragments used in this study**

#### 2. SUPPLEMENTAL FIGURES

**Figure S1. Activity assays performed with mature bacteriocins produced *in vitro* by a cell free system**

**Figure S2. Activity assays performed with single peptides of class IIb salivaricins produced *in vivo***

**Figure S3. Spot-on-lawn assays of HSISS4 derivatives producing PsnJ, PsnK, or PsnL against eight Gram-positive bacteria**

#### 3. SUPPLEMENTAL APPENDICES

**Appendix S1. Multiparametric bioinformatics script used to identify class II bacteriocins**

**Appendix S2. Raw and treated data obtained with the bioinformatics script on 100 *S. salivarius* genomes**

This file includes Table S1-S8 and Fig. S1-S2.

### 1. SUPPLEMENTAL TABLES

#### Table S1. ComR and BlpR binding sites used for *in silico* bacteriocin searches in streptococci.

| Species or specific strain | Regulator | Promoter | Nucleotide sequence (5’-3’)[a] |
|---|---|---|---|
| *S. salivarius/S. thermophilus* | ComR | | |
| | | $P_{comX}$ | ATAGTGACATATATGTCGCTATTTTATTT |
| | | $P_{comS}$ | GTGGTGACATAAATGTCACTACTTTTTTA |
| | | $P_{00176}$ | ATAGTGACATTTATGTCACTATTTTTATA |
| | | $P_{slvX}$ | ATAGTGACATTTATGTCACTATTTTTTAT |
| | | $P_{blpK}$ | GTAGTGACATTTATGTCACTACTTTTTTG |
| | | $P_{01651}$ | ATGGTGACATCTATGTCACTATTTTTTAT |
| | | $P_{slvW}$ | ATAGTGACATTTATGTCACTATTTTTTTG |
| | | $P_{01584}$ | GTAGTGACATTTATGTCACTACTTTTTTA |
| *S. vestibularis* NCTC12167 | ComR | | |
| | | $P_{comX}$ | ATAGT**GACA**TATA**TGTC**TCTATTTTATTT |
| | | $P_{comS}$ | GTAGT**GACA**TTTA**TGTC**ACTACTTTTTTT |
| *S. sobrinus* SL1 | ComR | | |
| | | $P_{comX}$ | ATAGT**GACA**TTGA**TGTC**ACTACCAAATTT |
| | | $P_{comS}$ | AAGGT**GACA**TAAA**TGTC**GTTTTGTTTTGA |
| *S. downei* NCTC11391 | ComR | | |
| | | $P_{comX}$ | ATAGT**GACA**TTTA**TGTC**ACTACAAATTAT |
| | | $P_{comS}$ | ATAGT**GACA**TGGA**TGTC**ACTATCTTTTTG |
| *S. thermophilus* LMD-9 | BlpR | | |
| | | $P_{blpA}$ | ACCGTTTGGGACGTATGGCAACTTTTTGGGACG |
| | | $P_{blpD}$ | ACCATTCGGGACGTATAGCCACTTTTTGGGACG |
| | | $P_{blpU}$ | ACTACTCGGGACATATAGCCACTTTTTGGGACG |
| | | $P_{blpG}$ | ACCATTCGGGACGTATAGTCACTTTTTGGGACG |
| | | $P_{blpE}$ | ACCATTCGGGACATATAGCCACTTTTTGGGACG |
| *S. pneumoniae* TIGR4 | BlpR | | |
| | | $P_{blpA}$ | CATTCAGGAAGTTTTAATGACTATTCAAGA |
| | | $P_{blpK}$ | AATTCAAGACGTTTCGATGCCAATTCAAGA |
| | | $P_{blpY}$ | TATACTAAACTTATCAAAGTTACAACAAGA |
| | | $P_{blpT}$ | AATTCAAGACATTTCAATGACAATTAAAGA |
| *S. oralis* FDAARGOS_885 | BlpR | | |
| | | $P_{blpA}$ | CATTCAGGAAGTTTCAATGACAATTCAAGA |
| | | $P_{blpM}$ | AATTCAAGACGTTTCGATGACAATTCAAGA |
| | | $P_{blpQ}$ | AGTTCAAGACGTTTCGATGACAATTCAAGA |
| | | $P_{blpT}$ | CATTCAGGAAATTTCAATGACAATTAAAGA |

[a] The sequences of ComR- or BlpR-binding sites were used by the bioinformatic script to generate a weight matrix for *in silico* bacteriocin searches. When only two ComR-binding sites are available, the following consensus sequence **GACA**NNNN**TGTC** was used.

**Table S2. Natural variants of class II salivaricins.**

| Name | Species | Freq.[a] | Id. (%)[b] | Mature sequence (aa)[c] |
|---|---|---|---|---|
| **Single-peptide bacteriocin** | | | | |
| BlpK | *S. salivarius* | 0.26 | - | GCSWGGFAKQGVATGVGNGLRLGIKTRTWQGAVAGAAGGAIVGGVGYGATCWW |
| | *S. thermophilus* | 0.42 | 98 | V.................................................... |
| | *S. thermophilus* | 0.33 | 100 | ..................................................... |
| | *S. pneumoniae* | 0.24 | 74 | ..N..D...A..GG.AVR..Q...........AT.......L...A.A..... |
| | *S. pneumoniae* | 0.15 | 70 | ..N..D...A..GGAAAR..Q.....G.....AT.......L...A.A..... |
| | *S. pneumoniae* | 0.12 | 72 | ..N..D...A..GG.AAR..Q...........AT..V....L...A.A..... |
| | *S. pneumoniae* | 0.08 | 72 | ..N..D...A..GG.AAR..Q.....G.....AT.......L...A.A..... |
| | *S. pneumoniae* | 0.07 | 74 | ..N..D...A..GG.AAR..Q...........AT.......L...A.A..... |
| | *S. pneumoniae* | 0.03 | 72 | ..N..D...A..GG.AVR..Q...........AT..V....L...A.A..... |
| | *S. pneumoniae* | 0.02 | 75 | ..N..D...A..GG..AR..Q...........AT.......L...A.A..... |
| | *S. pyogenes* | 0.26 | 74 | -...H.L.QAA.F..A...F...V........A......LVI........... |
| BlpE | *S. salivarius* | 0.11 | - | RVNWERWGMCGASVAVGA---------------------------TIGGGVALLGC |
| | *S. salivarius* | 0.04 | 48 | ..................SEGFSAAAGGTAFFIGPYAIGTGAVGAA.......... |
| | *S. salivarius* | 0.01 | 97 | .............................................A.......... |
| | *S. thermophilus* | 0.12 | 50 | ..................SEGFSAAAGGTAFFIGPYAIGTGAVGAA.......... |
| | *S. thermophilus* | 0.09 | 48 | ........V.........SEGFSAAAGGTAFFIGPYAIGTGAVGAA.......... |
| | *S. thermophilus* | 0.01 | 48 | ..................SEGFSAAAGGTAFFIGPYAIGTGAVGAA.........- |
| BlpF | *S. salivarius* | 0.03 | - | AVCLMPNVNNKEG---DPLKDGWVSPPRYRSGEAYPMVYLPVCAIM |
| | *S. salivarius* | 0.02 | 88 | ...R.....S.................K...........I.I.... |
| | *S. salivarius* | 0.01 | 98 | ..........................T................... |
| | *S. salivarius* | 0.01 | 67 | VS.I..RT...A.GSY..FI..R....R.............I.VL. |
| | *S. thermophilus* | 0.22 | 100 | .............................................. |
| | *S. thermophilus* | 0.09 | 66 | VS.I..RT...A.GSY..FI..R..................I.VL. |
| SlvV | *S. salivarius* | 0.53 | - | ACSFWGATAAVGVGAVGGAIKGGIQTGTWQGAALKGIGYGIKDGITYGLICRY |
| | *S. salivarius* | 0.10 | 94 | ...................V......................G......T... |
| | *S. salivarius* | 0.04 | 96 | ..........................................G......T... |
| | *S. salivarius* | 0.01 | 98 | V.................................................... |

| | | | | |
|---|---|---|---|---|
| | *S. salivarius* | 0.01 | 98 | ...............................................E..... |
|  | *S. salivarius* | 0.01 | 83 | ...................V....................--------..... |
| | *S. salivarius* | 0.01 | 89 | .....E.............V.....................NG.....FT... |
| | *S. thermophilus* | 0.10 | 62 | G.NWK..A.T.A..........AVT.YS.......AV.....A.VA..AT..W |
| SlvW | *S. salivarius* | 0.41 | - | GFVSKPQTLPERLGWNKWWLKRRPPYGD |
| | *S. salivarius* | 0.18 | 86 | D...R.................KY.... |
| | *S. salivarius* | 0.12 | 93 | D...R....................... |
| | *S. salivarius* | 0.12 | 82 | D...R...........R.....KY.... |
| | *S. salivarius* | 0.03 | 68 | N..YR.K.F......DR.....KY.... |
| | *S. salivarius* | 0.02 | 82 | D...R...........R.....KH.... |
| | *S. salivarius* | 0.02 | 79 | D...R.....G.....R.....KY.... |
| | *S. salivarius* | 0.01 | 82 | D...R....S............KY.... |
| | *S. salivarius* | 0.01 | 79 | D...R..........DR.....KY.... |
| | *S. salivarius* | 0.01 | 96 | ....R....................... |
| | *S. salivarius* | 0.01 | 75 | D...S....T......R....IKY.... |
| | *S. salivarius* | 0.01 | 89 | ...AR..................S.... |
| SlvX | *S. salivarius* | 0.11 | - | GCSWNNAAKAALGTSITGLITSGPGGALLGLAGGAVSYGALCWV |
| | *S. salivarius* | 0.01 | 91 | ..............N......NCS.................... |
| PsnI | *S. salivarius* | 0.09 | - | GVIGCVAGTAGSAGLGFLTGTSVGTVTFPIVGTVSGGAFGAWSGAGLGMATFCGV |
| | *S. salivarius* | 0.04 | 98 | ..........E............................................ |
| | *S. salivarius* | 0.02 | 96 | D.........................A............................ |
| | *S. salivarius* | 0.01 | 96 | .......V.............................V................. |
| | *S. salivarius* | 0.01 | 67 | .WVR.AL.............SGA..F.......I....I.G...GAV.I....-- |
| | *S. thermophilus* | 0.06 | 82 | ........A...PV.R.........H.L.....I...V........V...I.... |
| | *S. thermophilus* | 0.04 | 75 | ........AD..PV.R..P.I....H.L.....I...V........V...I.... |
| | *S. thermophilus* | 0.01 | 78 | ........AD..PV.R........SH.L.....I...V........V...I.... |
| PsnJ | *S. salivarius* | 0.13 | - | KVNWDRLSSSASGAAEGIYFCAASGAPLFTLAPYAIVGCGVVGAGLGYAFPH |
| PsnK | *S. salivarius* | 0.02 | - | KVIYYGNGLYCKSNGGCWVDWSQTINSILTNSAMNWATRGNAGWHSGGIAP |
| PsnL | *S. salivarius* | 0.50 | - | GWVKCYAGTIGSALVGSAGGPVGYWGGALVGYATFC |
| PsnM | *S. salivarius* | 0.01 | - | GCSWRGAGGATVQGAIGGA--------FGGNVVLPVVGSVPGYLAGGVLGGAGGAVAYGATCWWS |

| | | | | |
|---|---|---|---|---|
|  | *S. salivarius* | 0.01 | 80 | ..........AA.......IAGA----...............WVG..A...V..T.......... |
|  | *S. thermophilus* | 0.39 | 86 | ....G..............IGGA----...........................T.......... |
|  | *S. thermophilus* | 0.22 | 98 | ...................--------...........................T.......... |
| | *S. thermophilus* | 0.06 | 85 | ....G..............IGGAIGGA...........................T.......... |
|  | *S. thermophilus* | 0.04 | 96 | ......T............--------...........................T.......... |
|  | *S. thermophilus* | 0.04 | 80 | ......T...A........IAGA----...............WVG..A...V..T.......... |
|  | *S. thermophilus* | 0.03 | 80 | ..........AA.......IAGA----...............WVG..A...V..T.......... |
|  | *S. thermophilus* | 0.03 | 92 | ...................IGGA----...........................T.......... |
|  | *S. thermophilus* | 0.03 | 90 | ......T............IGGA----...........................T.......... |
|  | *S. thermophilus* | 0.01 | 89 | ......T............IGGA----...........................T..H....... |
| | *S. thermophilus* | 0.01 | 86 | ...................IGGAIGGA...........................T.......... |
| **Two-peptide bacteriocin** | | | | |
| SlvYZ | | | | |
| SlvY | *S. salivarius* | 0.46 | - | GWKTNLAIGGLCLASGPIGTMVCLGAYNGYMDSAR |
| | *S. salivarius* | 0.22 | 97 | .....................M............. |
| | *S. salivarius* | 0.16 | 97 | ...............R................... |
| | *S. salivarius* | 0.06 | 94 | .......V.............I............. |
| | *S. salivarius* | 0.04 | 97 | .....................I............. |
| | *S. salivarius* | 0.03 | 97 | .....I............................. |
| | *S. salivarius* | 0.02 | 89 | .....I..............L..V......K.... |
| | *S. salivarius* | 0.01 | 94 | .....................M........V.... |
| SlvZ | *S. salivarius* | 0.78 | - | VVPWAAISVGIAAAKLTYDLSYAAGKSFYNLTH |
| | *S. salivarius* | 0.05 | 94 | ....V.....M...................... |
| | *S. salivarius* | 0.04 | 97 | .I............................... |
| | *S. salivarius* | 0.04 | 97 | ...............................I. |
| | *S. salivarius* | 0.03 | 91 | ....V.....M...........T.......... |
| | *S. salivarius* | 0.02 | 91 | ....V.....M............T......... |
| | *S. salivarius* | 0.01 | 94 | ....G.....M...................... |
| | *S. salivarius* | 0.01 | 97 | ..........M...................... |
| | *S. salivarius* | 0.01 | 91 | A.........MV..................... |
| | *S. salivarius* | 0.01 | 85 | A.....V................V..TY..... |

| | | | | |
|---|---|---|---|---|
| | *S. thermophilus* | 0.55 | 88 | T...........G...............C..K. |
| | *S. thermophilus* | 0.06 | 85 | T.....V.....G...............C..K. |
| | *S. thermophilus* | 0.03 | 85 | T...........G...............CD.K. |
| | *S. thermophilus* | 0.01 | 85 | T.......A...G...............C..K. |
| | | | | PsnAB |
| PsnA | *S. salivarius* | 0.07 | - | GILSSVAGLVKDTWGTLYSTGRDFGRSVVNAAMP |
| | *S. salivarius* | 0.05 | 94 | ..M............................... |
| | *S. pyogenes* | 0.02 | 65 | .VV.DIG.F.GN..N...N........I...V.. |
| PsnB | *S. salivarius* | 0.07 | - | VAPVVIAVGVVATPFVLGAIAGGADEYARKRGHH |
| | *S. salivarius* | 0.05 | 97 | ..........I....................... |
| | *S. pyogenes* | 0.02 | 71 | .G.LFV..ALI...................K..R |
| | | | | PsnCD |
| PsnC | *S. salivarius* | 0.12 | - | NRCKDLIFGGALTGAGAGFTGGMAAFGVTAFPGAFVGAHVGAVAGGLACIGASF |
| | *S. salivarius* | 0.01 | 98 | .......................V.............................. |
| PsnD | *S. salivarius* | 0.12 | - | GKVGAVIGGCLGGMLMAWAAGPVSAGGYTMICATSGLANAYF |
| | *S. salivarius* | 0.01 | 98 | ....E..................................... |
| | | | | PsnEF |
| PsnE | *S. salivarius* | 0.11 | - | SIYSSVGYGVGVVTHTVYDFGRGFVDGFRG |
| PsnF | *S. salivarius* | 0.11 | - | NVAYDIGYGVGQVSYIAVEILKLRKIR |
| | *S. thermophilus* | 0.42 | 93 | ..P........P............... |
| | *S. thermophilus* | 0.35 | 89 | ..P.......EP............... |
| | *S. thermophilus* | 0.10 | 89 | ..P...V....P............... |
| | *S. thermophilus* | 0.08 | 89 | ..P.......RN............... |
| | *S. thermophilus* | 0.03 | 89 | ..P...S....P............... |
| | *S. thermophilus* | 0.01 | 89 | ..P........P....T.......... |
| | *S. thermophilus* | 0.01 | 89 | ..P.....E..P............... |
| | | | | PsnGH |
| PsnG | *S. salivarius* | 0.13 | - | GLAPIVIGGVVVGWKVIGGVAGVIAASGAAGIAAGYYANRP |
| PsnH | *S. salivarius* | 0.13 | - | VAPIMVAIGLGSLAVASFSSGYKFGTDLARRGR |

[a] Freq., frequency of the salivaricin among all analyzed strains of the species. The analysis was performed in *S. salivarius*. *S. thermophilus*, *S. pneumoniae*, and *S. pyogenes* from 99, 78, 137, and 261 genomes, respectively.
[b] Id. (%), percentage of identity for each salivaricin candidate compared to the prototypical bacteriocin reported on the top.
[c] Alignment of mature sequences of salivaricin variants. A dot or a minus sign indicate an identical or missing residue, respectively

#### Table S3. Distribution of class II bacteriocins in *S. salivarius* genomes.

| Strain | GenBank accession n° | SlvY | SlvZ | SlvW | SlvV | PsnL | BlpK | PsnI | BlpE | PsnJ | PsnC | PsnD | PsnG | PsnH | SlvX | PsnA | PsnB | PsnE | PsnF | BlpF | PsnK | PsnM | # Bac |
|---|---|---|---|---|---|---|---|---|---|---|---|---|---|---|---|---|---|---|---|---|---|---|---|
| **Circular genome** | | | | | | | | | | | | | | | | | | | | | | | |
| 57.I | GCA_000305335.2 | ■ | ■ | ■ | ■ | | ■ | | | | | | ■ | ■ | | | | | | | | | **7** |
| ATCC25975 | GCA_002094975.1 | ■ | ■ | ■ | | ■ | | ■ | | | ■ | ■ | | | | | | | | | | | **7** |
| ATCC27945 | GCA_002094955.1 | ■ | ■ | ■ | | | | | | | ■ | ■ | | | | ■ | ■ | | | | | | **7** |
| CCHSS3 | GCA_000253335.1 | ■ | ■ | ■ | ■ | | ■ | | | | | | ■ | ■ | | | | | | | | | **7** |
| DB-B5 | GCA_013822985.1 | ■ | ■ | ■ | ■ | | ■ | | | | | | | | | | | | | | | | **5** |
| FDAARGOS_259 | GCA_002073835.2 | ■ | ■ | ■ | ■ | ■ | | | | | | | | | | | | | | | | | **5** |
| FDAARGOS_771 | GCA_013267495.1 | ■ | ■ | ■ | ■ | | | | | ■ | | | ■ | ■ | | ■ | ■ | | | | | | **9** |
| FDAARGOS_1045 | GCA_016127535.1 | ■ | ■ | ■ | ■ | | | | | ■ | | | | | | | | | | | | | **5** |
| HSISS4 | GCA_000448685.2 | ■ | ■ | ■ | ■ | | ■ | | | | | | | | ■ | | | | | | | | **6** |
| ICDC1 | GCA_009738205.1 | ■ | ■ | ■ | ■ | ■ | | | | | | | | | | | | | | ■ | ■ | | **7** |
| ICDC2 | GCA_009738225.1 | ■ | ■ | ■ | ■ | | | | | ■ | | | | | | | | | | | | | **5** |
| ICDC3 | GCA_009738245.1 | ■ | ■ | ■ | ■ | ■ | | | | | | | | | | | | | | ■ | ■ | | **7** |
| JF | GCA_001543085.1 | ■ | ■ | ■ | ■ | | ■ | | ■ | | | | | | | | | | | | | | **6** |
| JIM8777 | GCA_000253315.1 | ■ | ■ | ■ | ■ | | ■ | | ■ | | | | | | | | | | | | | | **6** |
| LAB813 | GCA_008305695.1 | ■ | ■ | ■ | ■ | | ■ | | | | | | | | | | | | | | | | **5** |
| NCTC7366 | GCA_900475225.1 | ■ | ■ | ■ | | ■ | | ■ | | | ■ | ■ | | | | | | | | | | | **7** |
| NCTC8618 | GCA_900636435.1 | ■ | ■ | ■ | ■ | | ■ | | ■ | | | | | | | | | | | | | | **6** |
| SALI-10 | GCA_022936265.1 | ■ | ■ | | ■ | | ■ | | | | | | ■ | ■ | | | | | | | | | **6** |
| **Contig/scaffold**[a] | | | | | | | | | | | | | | | | | | | | | | | |
| 1001175st1_H3 | GCA_005844475.1 | ■ | ■ | ■ | ■ | | ■ | | ■ | | | | | | | | | | | | | | **6** |
| 1003_SOLI | GCA_001069855.1 | ■ | ■ | ■ | ■ | ■ | | | | | | | | | ■ | | | | | | | | **6** |
| 1270_SSAL | GCA_001069445.1 | ■ | ■ | ■ | ■ | ■ | | | ■ (magenta) | | | | | | | | | | | ■ (magenta) | | | **7** |
| 140_SSAL | GCA_001070595.1 | ■ | ■ | ■ (yellow) | ■ | ■ | | | | | | | ■ | ■ | | | | ■ | ■ | | | | **8** |
| 20-12 S2 | GCA_001659645.1 | ■ | ■ | ■ | | | | | | | ■ | ■ | | | | | | | | | | | **5** |
| 22-08 S7 | GCA_001906885.1 | ■ | ■ | ■ | ■ | ■ | | | | | | | | | | | | ■ | ■ | | | | **7** |
| 34-09 S8 | GCA_002093995.1 | ■ | ■ | ■ | | | | ■ | | ■ | | | | | | | | | | | | | **5** |
| 34-19 S9 | GCA_001906895.1 | ■ | ■ | ■ | | | | ■ | | ■ | | | | | | | | | | | | | **5** |
| 34-24 S10 | GCA_001906915.1 | ■ | ■ | ■ | | | | ■ | | ■ | | | | | | | | | | | | | **5** |
| 37-08 S12 | GCA_001906965.1 | ■ | ■ | ■ | ■ | | ■ | | | | | | | | | | | | | | | | **5** |
| 37-09 S13 | GCA_001906995.1 | ■ | ■ | ■ | ■ | ■ | | | | | | | | | | | | ■ | ■ | | | | **7** |
| 39-01 S14 | GCA_002094035.1 | ■ | ■ | ■ | | ■ | | | | | | | | | | | | ■ | ■ | | | | **6** |
| 39-07 S15 | GCA_001907045.1 | ■ | ■ | ■ | ■ | ■ | | | | | | | | | | ■ (magenta) | ■ (magenta) | | | | | | **7** |
| 39-09 S16 | GCA_001907005.1 | ■ | ■ | ■ | ■ | ■ | | | | | | | | | | ■ | ■ | | | | | | **7** |
| 40-02 S18 | GCA_001907055.1 | ■ | ■ | ■ | ■ | ■ | | | | | | | | | | | | ■ | ■ | | | | **7** |
| 726_SSAL | GCA_001074945.1 | ■ | ■ | ■ | ■ | ■ | | | | | ■ | ■ | | | | | | | | | | | **7** |
| 84-12 S20 | GCA_001906975.1 | ■ | ■ | ■ | | ■ | | | | | | | ■ | ■ | ■ | ■ | ■ | | | | | | **9** |
| 85-02 S21 | GCA_002094045.1 | ■ | ■ | ■ | | | | ■ | | ■ | | | | | | | | | | | | | **5** |
| 85-04 S22 | GCA_001907065.1 | ■ | ■ | ■ | | | | ■ | | ■ | | | | | | | | | | | | | **5** |
| 85-05 S23 | GCA_001907105.1 | ■ | ■ | ■ | | | | ■ | | ■ | | | | | | | | | | | | | **5** |
| 85-06 S24 | GCA_002094005.1 | ■ | ■ | ■ | | | | ■ | | ■ | | | | | | | | | | | | | **5** |
| AF10-23 | GCA_003465885.1 | ■ | ■ | ■ | ■ | | ■ | | | | | | | | | | | | | | | | **5** |
| AF23-9AC | GCA_003458645.1 | ■ | ■ | ■ | ■ | ■ | | | | | | | | | | | | | | | | | **5** |
| AF29-16 | GCA_003457695.1 | ■ | ■ | ■ | ■ | | ■ | | | | | | | | | | | | | | | | **5** |

| | | |
|---|---|---|
| ATCC 7073 | GCA_900143035.1 | **6** |
| BCC42 | GCA_003942515.1 | **7** |
| bf_0095 | GCA_004166915.1 | **6** |
| BIOML-A1 | GCA_009717495.1 | **10** |
| BIOML-A2 | GCA_009717465.1 | **10** |
| BIOML-A3 | GCA_009717475.1 | **6** |
| BIOML-A4 | GCA_009717435.1 | **6** |
| BIOML-A5 | GCA_009717425.1 | **6** |
| BIOML-A6 | GCA_009717395.1 | **9** |
| BIOML-A7 | GCA_009717375.1 | **6** |
| BIOML-A8 | GCA_009717365.1 | **9** |
| BIOML-A9 | GCA_009717335.1 | **9** |
| BIOML-A10 | GCA_009717325.1 | **6** |
| BIOML-A13 | GCA_009717265.1 | **5** |
| BIOML-A17 | GCA_009717205.1 | **5** |
| BIOML-A18 | GCA_009717155.1 | **5** |
| BIOML-A19 | GCA_009717165.1 | **5** |
| BIOML-A20 | GCA_009717145.1 | **5** |
| BIOML-A21 | GCA_009717105.1 | **5** |
| BIOML-A22 | GCA_009717115.1 | **5** |
| BIOML-A23 | GCA_009717065.1 | **5** |
| BIOML-A24 | GCA_009717045.1 | **5** |
| BIOML-A25 | GCA_009717055.1 | **5** |
| BIOML-A26 | GCA_009716965.1 | **5** |
| BIOML-A27 | GCA_009716945.1 | **5** |
| BIOML-A28 | GCA_009717005.1 | **5** |
| BIOML-A30 | GCA_009716955.1 | **5** |
| BIOML-A31 | GCA_009716865.1 | **5** |
| BIOML-A32 | GCA_009716845.1 | **5** |
| BIOML-A33 | GCA_009716875.1 | **5** |
| BIOML-A34 | GCA_009716785.1 | **5** |
| BIOML-A35 | GCA_009716765.1 | **5** |
| DE0578 | GCA_007665725.1 | **7** |
| F6-1 | GCA_002925865.1 | **7** |
| HS0302 | GCA_003231475.1 | **6** |
| K12 | GCA_000286295.1 | **5** |
| KB005 | GCA_000963335.1 | **5** |
| M18 | GCA_000225385.2 | **5** |
| MGYG-HGUT-00113 | GCA_902363415.1 | **5** |
| MIT 14-1770-C1 | GCA_002417605.1 | **5** |
| NU10 | GCA_000714445.1 | **8** |
| Ssal B50) | GCA_902850315.1 | **5** |
| Ssal B63 | GCA_902850305.1 | **5** |
| Ssal B57 | GCA_902850285.1 | **7** |
| Ssal L22 | GCA_902858925.1 | **6** |
| Ssal L25 | GCA_902858935.1 | **6** |
| Ssal L45 | GCA_902858945.1 | **7** |
| Ssal L61 | GCA_902859045.1 | **8** |

| | | | | | | | | | | | | | | | | | | | | | | |
|---|---|---|---|---|---|---|---|---|---|---|---|---|---|---|---|---|---|---|---|---|---|---|
| Ssal L64 | GCA_902859025.1 | | | | | | | | | | | | | | | | | | | | | **5** |
| Ssal N20 | GCA_902859005.1 | | | | | | | | | | | | | | | | | | | | | **6** |
| Ssal N5 | GCA_902859015.1 | | | | | | | | | | | | | | | | | | | | | **5** |
| PS4 | GCA_000257585.1 | | | | | | | | | | | | | | | | | | | | | **6** |
| UBSS01 | GCA_006227745.1 | | | | | | | | | | | | | | | | | | | | | **10** |
| UC3162 | GCA_000963265.1 | | | | | | | | | | | | | | | | | | | | | **5** |
| UMB0051 | GCA_002860785.1 | | | | | | | | | | | | | | | | | | | | | **5** |
| VA08-2AN | GCA_011029085.1 | | | | | | | | | | | | | | | | | | | | | **7** |
| YU10 | GCA_000714455.1 | | | | | | | | | | | | | | | | | | | | | **8** |
| | **Number of hit** | **99** | **99** | **94** | **70** | **50** | **26** | **17** | **16** | **13** | **13** | **13** | **13** | **13** | **11** | **12** | **12** | **10** | **10** | **7** | **2** | **2** |

[a] A colored box indicates that the bacteriocin have been identified. Grey boxes represent bacteriocins found through the bioinformatic screen. Bacteriocins represented by boxes of other colors were not identified by the script but were discovered using tBLASTn. Red boxes are bacteriocins without an upstream ComR-box. Purples boxes are bacteriocins at the start/end of a contig, preventing to identify the associated ComR-box. Yellow boxes indicate truncated bacteriocins.

**Table S4. Bacteriocin candidates identified by *in silico* analysis in other streptococci.**

| Name | Precursor sequence (aa)[a] | MW (kDa) | Gly-Ala (%)[b] | Lys-Arg (%)[c] | GRAVY[d] | pI[e] | Charge[f] |
|---|---|---|---|---|---|---|---|
| **ComRS system** | | | | | | | |
| *Streptococcus vestibularis* NCTC 12167 | | | | | | | |
| **SlvY** | MINKEMKAAE**L**ASVT**GG**↓GWKTNLAVGVGGLCLASGPIGTLICIGSYNGYMDTAR | 3.7 | 28.9 | 5.2 | 0.5 | 8.2 | 1 |
| **SlvW** | MRTKAYGEELNAET**L**ENVT**GG**↓GFVARPQTLPERLGWNKWWLKRSPPYGD | 3.4 | 14.3 | 17.9 | -1 | 10.9 | 3 |
| **PsnE** | MKNAFQKIDIKD**L**VIVV**GG**↓SIDSSVGYGVGVVTHTVYDFGRGFVDGFRG | 3.0 | 23.3 | 6.7 | 0.3 | 5.5 | -0.5 |
| **PsnF** | MDTMMMDKFDSLTFDK**L**SEIV**GG**↓NVAYDIGYGVGQVSYIAVEILKLRKIR | 3.0 | 18.5 | 14.8 | 0.3 | 9.7 | 2 |
| *Streptococcus downei* NCTC 11391 | | | | | | | |
| **Bac1** | MDTLALDNFEAANMSD**L**SEVQ**GG**↓TNPTRCVVGIAGEGVQSGIAGAAALGAVGLAVGG PVGAYGGAHLGFFLGFIGGSLKGTADYCF | 5.9 | 42.9 | 3.2 | 0.8 | 7.3 | 0.5 |
| **Bac2** | MNTVALDNFEAANMSE**L**SEVQ**GG**↓FSWQDAISCSCCNTISPFRCSSWCLCIMGNYKTTG HIKLEIIFFR | 5.2 | 6.7 | 8.9 | 0.3 | 8.1 | 2.5 |
| **Bac3** | MAFDNFEVTNAKF**L**AESV**GG**↓VNEVVAGSACAMALGFVGTAILAGSGPVGWATLALA YACDGVAIAALNDK | 4.8 | 40.0 | 2.0 | 1.1 | 3.9 | -2 |
| **Bac4** | MDTKAFDTFEVVEDDQ**L**SDTT**GG**↓YSKYDITISGITALGYGFAAFGALACPPAALAFVAA EGIQGAITAYMAYR | 5.1 | 37.2 | 3.9 | 0.8 | 6.3 | 0 |
| **Bac5** | MNKDVLSNFETLDNTQ**L**VKIK**GG**↓GDEGYNFWYGIGKWARQQYRGFCNWSNWCKG | 3.8 | 21.9 | 12.5 | -1.1 | 3.9 | 2 |
| *Streptococcus sobrinus* SL1 | | | | | | | |
| **Bac1** | MNYVELTTDE**L**AAIE**GG**↓NILHSIGNAFSSAYHGLVDGWNNH | 2.6 | 20.0 | 0.0 | -0.2 | 6.8 | 0.5 |
| **Bac2** | METKTFEKYETVNVEE**L**AELV**GG**↓NTAYDAGKVVGKVGQIAAVIAAFF | 2.4 | 36.0 | 8 | 0.9 | 9.3 | 1 |
| **Bac3** | MDTLALDNFEAANMSE**L**SEVQ**GG**↓FSWQDAIPVAIATLYPPLGAALGVYALWGTIKPQG ILS | 4.0 | 25.6 | 2.6 | 0.8 | 6.3 | 0 |
| **Bac4** | MKTKELLFSELNEED**L**AAIR**GG**↓SLWDFKGIIIGDSWWYPRIGIPEHPILLDK | 3.6 | 9.7 | 9.7 | 0.0 | 5.6 | -0.5 |

| | | | | | | | |
|---|---|---|---|---|---|---|---|
| **Bac5** | MTTFRSCQRHWRLLF**V**VMEK**GG**↓NRR**C**WL**C**LWTNVLNQMFLLIWSLKLVSSSFFKFMKNWG | 4.7 | 2.6 | 12.8 | 0.35 | 11.1 | 5 |
| **Bac6** | MNKDVLSNFETLDNTQ**L**VKIK**GG**↓GDEGYNFWYGIGKWARQQYRGF**C**NWSNW**C**KG | 3.8 | 21.9 | 12.5 | -1.1 | 8.8 | 2 |
| **Bac7** | MIIWRGKGLLL**L**VSIFA**GG**↓IASGLVGSMAANLSVGPLKSLVAFLAALAFGVSALVNHIFCKTLLKNEG | 4.9 | 26.0 | 6 | 1.1 | 9.6 | 2.5 |
| **BlpRH system** | | | | | | | |
| *Streptococcus pneumoniae* TIGR4 | | | | | | | |
| **BlpI** | MNTKMMEQFSVMDNEE**L**EIVS**GG**↓RGNLGSAIGG**C**IGAVLLAAATGPITGGAATLI**C**VGSGIMSSL | 3.8 | 40.5 | 2.4 | 1.2 | 8.2 | 1 |
| **BlpJ** | MNTKMLSQLEVMDTEM**L**AKVE**GG**↓YSSTD**C**QNALITGVTTGIITGGTGAGLATLGVAGLAGAFVGAHIGAIGGGLT**C**LGGMVGDKLGLSW | 6.1 | 39.4 | 1.5 | 0.8 | 5.4 | -0.5 |
| **BlpK** | MDTKMMSQFSVMDTEM**L**ACVE**GG**↓G**C**NWGDFAKAGVGGGAARGLQLGIKTGTWQGAATGAAGGAILGGVAYAAT**C**WW | 5.1 | 50.9 | 5.7 | 0.3 | 8.8 | 2 |
| **BlpM** | MDTKIMEQFHEMDITM**L**SSIE**GG**↓KNNWQTNVLEGGGAAFGGWGLGTAI**C**AASGVGAPFMGA**C**GYIGAKFGVDLWAGVTGATGGF | 5.9 | 44.3 | 3.3 | 0.5 | 6.2 | 0 |
| **BlpN** | MNTYCNINETM**L**SEVY**GG**↓NSGGAAVVAALG**C**AAGGVKYGRLLGPWGAAIGGIGGAVV**C**GYLAYTATS | 4.5 | 49.0 | 4.1 | 0.9 | 8.8 | 2 |
| **BlpO** | MDTKMMSQFAVMDNEM**L**ACVE**GG**↓DIDWGRKIS**C**AAGVAYGAIDG**C**ATTV | 2.6 | 34.6 | 7.7 | 0.4 | 4.3 | -1 |
| *Streptococcus oralis* FDAARGOS_885 | | | | | | | |
| **BlpD** | MNTKMMEQFKIMDTEM**L**ASIE**GG**↓TDWGTVGKGAVYGAGIGVAM**C**TVGGLLGGSAWAMTAG**C**AWAGAKLGGAFTAIADNIWP | 5.7 | 44.1 | 3.4 | 0.7 | 5.6 | 0 |
| **BlpE** | MFDYKIVDNQE**L**SNIS**GG**↓GLGGDVVVGALSGAFQAGQS**C**IAGGPQAYLI**C**ATGGAIVGGILAFGLRPPK | 4.8 | 41.2 | 3.9 | 0.8 | 4.8 | 1 |
| **BlpN** | MNTYYNVDETM**L**SEIY**GG**↓NSGGAAVVAALG**C**AAGGVKYGKFLGPWGAAIGGIGGALI**C**GYLAYSATS | 4.5 | 49.0 | 4.1 | 0.9 | 4.5 | 2 |
| **BlpM** | MDTKMIEQFHEMDITM**L**SSIE**GG**↓KNNWQTNVLEGGGAAFGGWGLGTAI**C**AASGVGAPFMGA**C**GYIGAKFGVALWAGVTGATGGF | 5.8 | 45.9 | 3.3 | 0.6 | 5.8 | 1 |

| | | | | | | | |
|---|---|---|---|---|---|---|---|
| **BlpQ** | MNTKMMSQFSVIDNEM**L**DRIE**GG**↓IFGVDDAVFWTVGGYVVGRIVDTAIGDFTNQ**<u>C</u>**RK NPHQWF**<u>C</u>**VRV | 5.0 | 15.9 | 9.1 | 0.2 | 5.0 | 0.5 |
| **PncT** | MKKIDYIALNEVE**L**ETIS**GG**↓DD**<u>C</u>**FIGDIG**<u>C</u>**IGWGILKSIGGMIKPGPYVPPV**<u>C</u>**IPKSSWNP APPVP**<u>C</u>** | 4.9 | 17.0 | 6.4 | 0.3 | 4.94 | 0 |
| **Bac1** | MNTKMMEQFESMDTDM**L**ACVE**GG**↓KKFGD**<u>C</u>**ETAISAGIGVGAVFAGPWGAVGLGTVT NMFF**<u>C</u>**ATPVS | 4.2 | 32.6 | 4.7 | 0.8 | 4.2 | 0 |
| **Bac2** | MNTKMMEQFEIMDTDM**L**AKVE**GG**↓FGGWGDMIAGLLGGLAPSPTLDQLNGKWPIIHFS KP**<u>C</u>**GPYGIGGTPNS**<u>C</u>**NGI | 5.3 | 26.9 | 3.8 | 0.1 | 5.3 | 0.5 |
| **Bac3** | MNLKMMEQFEIMDTEM**L**ASKV**GG**↓ KTIYYGNGLY**<u>C</u>**DNSKG**<u>C</u>**WVNWPEAINKILTNSIVNGFSGGNAGWNSGGPL | 5.4 | 22.0 | 6.0 | -0.3 | 5.4 | 1 |
| *Streptococcus thermophilus* LMD-9 | | | | | | | |
| **BlpD** | MATQTIENFNTLDLET**L**ASVE**GG**↓LS**<u>C</u>**DEGMLAVGGLGAVGGPWGAVGGVLVGAALY **<u>C</u>**F | 3.3 | 42.9 | 0.0 | 1.2 | 3.6 | -2 |
| **BlpE** | MATQTIENFNTLDLET**L**ASVE**GG**↓RVNWERWGM**<u>C</u>**GASVAVGASEGFSAAAGGTAFFIG PYAIGTGAVGAAIGGGVALLG**<u>C</u>** | 5.3 | 46.4 | 3.6 | 0.8 | 6.3 | 0 |
| **BlpF** | MFTKLTHND**L**EVIQ**GG**↓AV**<u>C</u>**LMPNVNNKEGDPLKDGWVSPPRYRSGEAYPMVYLPV**<u>C</u>** AIM | 4.8 | 14.0 | 9.3 | -0.1 | 6.4 | 0 |
| **BlpK** | MATQTIENFNTLDLET**L**ASVE**GG**↓G**<u>C</u>**SWGGFAKQGVATGVGNGLRLGIKTRTWQGAVA GAAGGAIVGGVGYGAT**<u>C</u>**WW | 5.1 | 45.3 | 7.5 | 0.3 | 10.3 | 4 |

[a] The predicted cleavage motif ([**M**|**L**|**V**]$X_4$**GG**↓) of the leader sequence is indicated. The cysteine residues potentially involved in disulfide bridges are bold underlined.
[b] % Gly-Ala, percentage in glycine and alanine in the mature sequence.
[c] % Lys-Arg, percentage in lysine and arginine in the mature sequence.
[d] GRAVY, GRand AVerage of hydropathY of the mature sequence, positive and negative values indicate hydrophobic and hydrophilic peptides, respectively.
[e] pI, isoelectric point of the mature sequence.
[f] Charge, charge of the bacteriocin (D and E [-1], H [+0.5], K and R [+1]).

**Table S5. List of bacterial strains used in this study.**

| Names | Characteristics | Reference/source |
|---|---|---|
| *Streptococcus salivarius* | | |
| HSISS4 | Wild-type gastro-intestinal tract isolate | (5) |
| *Δslv5* | HSISS4 Δ*slvX-HSISS4_01664*::*lox72* Δ*blpKl*::*lox72* Δ*slvY-HSISS4_01744* ::*lox72* Δ*slvV* ::*lox72* Δ*slvW-blpG* ::*lox72* | (3) |
| JM1015 | HSISS4 $tRNA^{ser}$::$P_{xyl1}$-*comR-spec* | (3) |
| JD0001 | *Δslv5* $tRNA^{Ser}$ ::$P_{blpK}$-01585-*spec* | This work |
| JD0002 | *Δslv5* $tRNA^{Ser}$ ::$P_{blpK}$-*blpE-spec* | This work |
| JD0003 | *Δslv5* $tRNA^{Ser}$ ::$P_{blpK}$-*blpF-spec* | This work |
| JD0004 | *Δslv5* $tRNA^{Ser}$ ::$P_{blpK}$-*slvV-spec* | This work |
| JD0005 | *Δslv5* $tRNA^{Ser}$ ::$P_{blpK}$-*slvW-spec* | This work |
| JD0006 | *Δslv5* $tRNA^{Ser}$ ::$P_{blpK}$-*slvX-spec* | This work |
| JD0007 | *Δslv5* $tRNA^{Ser}$ ::$P_{blpK}$-*slvY-spec* | This work |
| JD0008 | *Δslv5* $tRNA^{Ser}$ ::$P_{blpK}$-*slvZ-spec* | This work |
| JD0009 | *Δslv5* $tRNA^{Ser}$ ::$P_{blpK}$-*psnA-spec* | This work |
| JD0010 | *Δslv5* $tRNA^{Ser}$ ::$P_{blpK}$-*psnB-spec* | This work |
| JD0011 | *Δslv5* $tRNA^{Ser}$ ::$P_{blpK}$-*psnC-spec* | This work |
| JD0012 | *Δslv5* $tRNA^{Ser}$ ::$P_{blpK}$-*psnD-spec* | This work |
| JD0013 | *Δslv5* $tRNA^{Ser}$ ::$P_{blpK}$-*psnE-spec* | This work |
| JD0014 | *Δslv5* $tRNA^{Ser}$ ::$P_{blpK}$-*psnF-spec* | This work |
| JD0015 | *Δslv5* $tRNA^{Ser}$ ::$P_{blpK}$-*psnG-spec* | This work |
| JD0016 | *Δslv5* $tRNA^{Ser}$ ::$P_{blpK}$-*psnH -spec* | This work |
| JD0017 | *Δslv5* $tRNA^{Ser}$ ::$P_{blpK}$-*psnI-spec* | This work |
| JD0018 | *Δslv5* $tRNA^{Ser}$ ::$P_{blpK}$-*psnJ-spec* | This work |
| JD0019 | *Δslv5* $tRNA^{Ser}$ ::$P_{blpK}$-*pnsK-spec* | This work |
| JD0020 | *Δslv5* $tRNA^{Ser}$ ::$P_{blpK}$-*psnL-spec* | This work |
| JD0021 | *Δslv5* $tRNA^{Ser}$ ::$P_{blpK}$-*psnM-spec* | This work |
| JD0022 | HSISS4 $tRNA^{Ser}$ ::$P_{blpK}$-*psnJ-spec* | This work |
| JD0023 | HSISS4 $tRNA^{Ser}$ ::$P_{blpK}$-*pnsK-spec* | This work |
| JD0024 | HSISS4 $tRNA^{Ser}$ ::$P_{blpK}$-*psnL-spec* | This work |
| JD0025 | HSISS4 $tRNA^{Ser}$ ::$P_{blpK}$-*psnM-spec* | This work |
| *Streptococcus thermophilus* | | |
| LMD-9 | Wild-type milk product isolate | ATCC, American Type Culture Collection, Rockville, MD |
| *Streptococcus vestibularis* | | |
| F0396 | Wild type | Craig Venter Institute, Rockville, MD |
| *Streptococcus oralis* | | |
| Si 0464 | Wild type | J. Mahillon, laboratory collection |
| *Streptococcus mitis* | | |
| LMG 14557 | Wild type | BCCM/LMG collection, Laboratory of Microbiology, Ghent University, Belgium |
| *Streptococcus pyogenes* | | |
| 4549 | Wild type | (4) |
| *Lactococcus lactis* | | |
| IL1403 | Laboratory strain | (1) |
| *Staphylococcus aureus* | | |
| ATCC 6538 | Wild type | ATCC, American Type Culture Collection, Rockville, MD |
| *Staphylococcus epidermidis* | | |
| LMG10273 | Wild type | BCCM/LMG collection, Laboratory of Microbiology, Ghent University, Belgium |

| | | |
|---|---|---|
| *Enterococcus faecium* | | |
| ATCC 19434 | Wild type | ATCC, American Type Culture Collection, Rockville, MD |
| *Enterococcus faecalis* | | |
| Si0159 | Wild type | J. Mahillon, laboratory collection |
| *Listeria monocytogenes* | | |
| ATCC 51777 | Wild type | ATCC, American Type Culture Collection, Rockville, MD |

**Table S6. List of plasmids used in this study.**

| Names | Characteristics | Reference/source |
|---|---|---|
| pUC57-BlpK | Plasmid containing the mature sequence of the BlpK. This plasmid is compatible with the kit PURExpress. | This work |
| pUC57-PsnA | Plasmid containing the mature sequence of the PsnA. This plasmid is compatible with the kit PURExpress. | This work |
| pUC57-PsnB | Plasmid containing the mature sequence of the PsnB. This plasmid is compatible with the kit PURExpress. | This work |
| pUC57-PsnC | Plasmid containing the mature sequence of the PsnC. This plasmid is compatible with the kit PURExpress. | This work |
| pUC57-PsnD | Plasmid containing the mature sequence of the PsnD. This plasmid is compatible with the kit PURExpress. | This work |
| pUC57-PsnE | Plasmid containing the mature sequence of the PsnE. This plasmid is compatible with the kit PURExpress. | This work |
| pUC57-PsnF | Plasmid containing the mature sequence of the PsnF. This plasmid is compatible with the kit PURExpress. | This work |
| pUC57-PsnG | Plasmid containing the mature sequence of the PsnG. This plasmid is compatible with the kit PURExpress. | This work |
| pUC57-PsnH | Plasmid containing the mature sequence of the PsnH. This plasmid is compatible with the kit PURExpress. | This work |
| pUC57-PsnI | Plasmid containing the mature sequence of the PsnI. This plasmid is compatible with the kit PURExpress. | This work |
| pUC57-PsnJ | Plasmid containing the mature sequence of the PsnJ. This plasmid is compatible with the kit PURExpress. | This work |
| pUC57-PsnK | Plasmid containing the mature sequence of the PsnK. This plasmid is compatible with the kit PURExpress. | This work |
| pUC57-PsnL | Plasmid containing the mature sequence of the PsnL. This plasmid is compatible with the kit PURExpress. | This work |
| pUC57-PsnM | Plasmid containing the mature sequence of the PsnM. This plasmid is compatible with the kit PURExpress. | This work |
| pGILF*spec* | pG+host9 derivative containing the spectinomycin resistance cassette P*spec-spec* downstream of *luxAB* | (2) |

**Table S7. List of oligonucleotides used in this study.**

| Names | Sequences |
|---|---|
| F_slvV (blpK leader) | 5'-TTGCAAACGTTGAAGGTGGTGCATGTAGTTTTTGGGGAGC-3' |
| R_slvV (spec) | 5'-TTTATTGGCCGGCCTTATTACTAATAACGACAAATAAGTCC-3' |
| F_slvW (leader BlpK) | 5'-GGCGCTTGCAAACGTTGAAGGTGGTGATTTTGTATCTAGACCTCA-3' |
| R_slvW (spec) | 5'-TTTATTGGCCGGCCTTATTACTAATCGCCATAAGG-3' |
| F_slvX (leader BlpK) | 5'-GGCGCTTGCAAACGTTGAAGGTGGTGGATGTAGTTGGAACAATGC-3' |
| R_slvX (spec) | 5'-TTTATTGGCCGGCCTTATTACAATCATACCCAACACAAC-3' |
| F_slvY (leader BlpK) | 5'-GGCGCTTGCAAACGTTGAAGGTGGTGGATGGAAGACTAACCTTGC-3' |
| R_slvY_spec | 5'-TTTATTGGCCGGCCTTATTATTATCGCGCAGAGTCC-3' |
| F_slvZ (leader BlpK) | 5'-GGCGCTTGCAAACGTTGAAGGTGGTGTAGTTCCTTGGGCCGCTAT-3' |
| R_slvZ (spec) | 5'-TTTATTGGCCGGCCTTATTATTAGTGGGTGAGGTTATAG-3' |
| F_blpF (blpK leader) | 5'-TTGCAAACGTTGAAGGTGGTGCTGTCTGTCGTATGCCAAA-3' |
| R_blpF (spec) | 5'-TTTATTGGCCGGCCTTATTATTACATAATAGCACAGACAGGTAAG-3' |
| F_blpE (leader BlpK) | 5'-GGCGCTTGCAAACGTTGAAGGTGGTCGAGTCAATTGGGAACGATG-3' |
| R_blpE (spec) | 5'-GGATCTTTATTGGCCGGCCTTATTATTAACAGCCAAGTAAGGCTAC-3' |
| F_PblpK (tRNAser) | 5'-CCATCTTCTTTTTAATTACTAAGATAATCTAGACCATAGC-3' |
| R_blpK_spec | 5'-TTTATTGGCCGGCCTTATTATCACCACCAGCATGTTGCT-3' |
| R_01585_spec | 5'- TTTATTGGCCGGCCTTATTATTAAGATTTCTTTTTAACGG-3' |
| F_psnA (blpK leader) | 5'-TTGCAAACGTTGAAGGTGGTGGAATTCTGTCAAGTGTAGC-3' |
| R_psnA (spec) | 5'-TTTATTGGCCGGCCTTATTATTAAGGCATAGCAGCGTTAAC-3' |
| F_psnB (blpK leader) | 5'-TTGCAAACGTTGAAGGTGGTGTAGCCCCTGTAGTTATTG-3' |
| R_psnB (spec) | 5'-TTTATTGGCCGGCCTTATTATCAGTGATGCCCTCTCTTTC-3' |
| F_psnC (blpK leader) | 5'-TTGCAAACGTTGAAGGTGGTAATAGATGTAAAGATTTAATC-3' |
| R_psnC (spec) | 5'-TTTATTGGCCGGCCTTATTACTAAAAGCTCGCTCCGATAC-3' |
| F_psnD (blpK leader) | 5'-TTGCAAACGTTGAAGGTGGTGGGAAAGTTGGAGAAGTAATTG-3' |
| R_psnD (spec) | 5'-TTTATTGGCCGGCCTTATTATCAAAAATAAGCATTTGCAAGC-3' |
| F_psnE (blpK leader) | 5'-TTGCAAACGTTGAAGGTGGTAGTATTTACTCATCTGTAGG-3' |
| R_psnE (spec) | 5'-TTTATTGGCCGGCCTTATTATTAACCTCTAAATCCATCTAC-3' |
| F_psnF (blpK leader) | 5'-TTGCAAACGTTGAAGGTGGTAATGTAGCATATGACATTGG-3' |
| R_psnF (spec) | 5'-TTTATTGGCCGGCCTTATTACTATCTTATTTTTCTCAGTT-3' |
| F_psnG (blpK leader) | 5'-TTGCAAACGTTGAAGGTGGTCTTGCACCGATTGTAATTGG-3' |
| R_psnG (spec) | 5'-TTTATTGGCCGGCCTTATTATTAAGGACGATTAGCATAAT-3' |

| | |
|---|---|
| F_psnH (blpK leader) | 5’-TTGCAAACGTTGAAGGTGGTGTTGCACCAATTATGGTTGC-3’ |
| R_psnH (spec) | 5’-TTTATTGGCCGGCCTTATTACTATCGTCCACGTCTAGCTA-3’ |
| F_psnI (blpK leader) | 5’-TTGCAAACGTTGAAGGTGGTGGTGTGATTGGTTGTGTAGC-3’ |
| R_psnI (spec) | 5’-TTTATTGGCCGGCCTTATTACTAAACTCCGCAAAATGTAG-3’ |
| F_psnJ (blpK leader) | 5’-TTGCAAACGTTGAAGGTGGTAAAGTCAATTGGGACCGCTT-3’ |
| R_psnJ (spec) | 5’-TTTATTGGCCGGCCTTATTATTAATGTGGGAAAGCATAAC-3’ |
| F_psnK (blpK leader) | 5’-TTGCAAACGTTGAAGGTGGTAAAGTCATTTATTACGGTAATG-3’ |
| R_psnK (spec) | 5’-TTTATTGGCCGGCCTTATTATTATGGAGCAATACCACCTG-3’ |
| F_psnL (blpK leader) | 5’-TTGCAAACGTTGAAGGTGGTGGATGGGTTAAGTGTTATGC-3’ |
| R_psnL (spec) | 5’-TTTATTGGCCGGCCTTATTATTAGCAGAATGTAGCATAGC-3’ |
| F_psnM (blpK leader) | 5’-TTGCAAACGTTGAAGGTGGTGGATGTAGTTGGAGAGGCGC-3’ |
| F_psnM (Spec) | 5’-TTTATTGGCCGGCCTTATTATTAGCTCCACCAGCAGGTTG-3’ |
| R_blpk_leader | 5’-ACCACCTTCAACGTTTGCAA-3’ |
| UR_tRNAser | 5’-AGTAATTAAAAAGAAGATGG-3’ |
| UF_tRNAser | 5’-CAAGATTAACCATGACCTTC-3’ |
| DF_tRNAser | 5’-TACCTAAAAAGTGTCCCTTC-3’ |
| DR_tRNAser2 | 5’-TTGGATAAGGTCTTGACTTC-3’ |
| F_spec | 5’-TAATAAGGCCGGCCAATAAA-3’ |
| R-spec | 5’-ATAGGATGAGAACTCCCATG-3’ |

**Table S8. List of PCR fragments used in this study.**

| Strain | PCR fragment | Template DNA | Primer 1 | Primer 2 |
|---|---|---|---|---|
| JD0001 | UP HR of *tRNA*$_{ser}$ | HSISS4 | UF_tRNAser | UR_tRNAser |
| | Spec fused-Dw HR of tRNA$_{ser}$ | JM1015 | F_spec | DR_tRNAser |
| | PblpK-01585 | HSISS4 | F_PblpK (tRNAser) | R_01585_spec |
| JD0002 | UP HR of *tRNA*$_{ser}$ – *blpK* leader | JD0001 | UF_tRNAser | R_blpk_leader |
| | Spec fused-Dw HR of tRNA$_{ser}$ | JM1015 | F_spec | DR_tRNAser |
| | Mature sequence *blpE* | pUC57-BlpE | F_blpE (blpK leader) | R_blpE (spec) |
| JD0003 | UP HR of *tRNA*$_{ser}$ – *blpK* leader | JD0001 | UF_tRNAser | R_blpk_leader |
| | Spec fused-Dw HR of tRNA$_{ser}$ | JM1015 | F_spec | DR_tRNAser |
| | Mature sequence *blpF* | pUC57-BlpF | F_blpF (blpK leader) | R_blpF (spec) |
| JD0004 | UP HR of *tRNA*$_{ser}$ – *blpK* leader | JD0001 | UF_tRNAser | R_blpk_leader |
| | Spec fused-Dw HR of tRNA$_{ser}$ | JM1015 | F_spec | DR_tRNAser |
| | Mature sequence *slvV* | HSISS4 | F_slvV (blpK leader) | R_slvV (spec) |
| JD0005 | UP HR of *tRNA*$_{ser}$ – *blpK* leader | JD0001 | UF_tRNAser | R_blpk_leader |
| | Spec fused-Dw HR of tRNA$_{ser}$ | JM1015 | F_spec | DR_tRNAser |
| | Mature sequence *slvW* | HSISS4 | F_slvW (blpK leader) | R_slvW (spec) |
| JD0006 | UP HR of *tRNA*$_{ser}$ – *blpK* leader | JD0001 | UF_tRNAser | R_blpk_leader |
| | Spec fused-Dw HR of tRNA$_{ser}$ | JM1015 | F_spec | DR_tRNAser |
| | Mature sequence *slvX* | HSISS4 | F_slvX (blpK leader) | R_slvX (spec) |
| JD0007 | UP HR of *tRNA*$_{ser}$ – *blpK* leader | JD0001 | UF_tRNAser | R_blpk_leader |
| | Spec fused-Dw HR of tRNA$_{ser}$ | JM1015 | F_spec | DR_tRNAser |
| | Mature sequence *slvY* | HSISS4 | F_slvY (blpK leader) | R_slvY (spec) |
| JD0008 | UP HR of *tRNA*$_{ser}$ – *blpK* leader | JD0001 | UF_tRNAser | R_blpk_leader |
| | Spec fused-Dw HR of tRNA$_{ser}$ | JM1015 | F_spec | DR_tRNAser |
| | Mature sequence *slvZ* | HSISS4 | F_slvZ (blpK leader) | R_slvZ (spec) |
| JD0009 | UP HR of *tRNA*$_{ser}$ – *blpK* leader | JD0001 | UF_tRNAser | R_blpk_leader |
| | Spec fused-Dw HR of tRNA$_{ser}$ | JM1015 | F_spec | DR_tRNAser |
| | Mature sequence *psnA* | pUC57-PsnA | F_psnA (blpK leader) | R_psnA (spec) |
| JD0010 | UP HR of *tRNA*$_{ser}$ – *blpK* leader | JD0001 | UF_tRNAser | R_blpk_leader |
| | Spec fused-Dw HR of tRNA$_{ser}$ | JM1015 | F_spec | DR_tRNAser |
| | Mature sequence *psnB* | pUC57-PsnB | F_psnB (blpK leader) | R_psnB (spec) |

| | | | | |
|---|---|---|---|---|
| JD0011 | UP HR of *tRNA*$_{ser}$ – *blpK* leader | JD0001 | UF_tRNAser | R_blpk_leader |
| | Spec fused-Dw HR of tRNA$_{ser}$ | JM1015 | F_spec | DR_tRNAser |
| | Mature sequence *psnC* | pUC57-PsnC | F_psnC (blpK leader) | R_psnC (spec) |
| JD0012 | UP HR of *tRNA*$_{ser}$ – *blpK* leader | JD0001 | UF_tRNAser | R_blpk_leader |
| | Spec fused-Dw HR of tRNA$_{ser}$ | JM1015 | F_spec | DR_tRNAser |
| | Mature sequence *psnD* | pUC57-PsnD | F_psnD (blpK leader) | R_psnD (spec) |
| JD0013 | UP HR of *tRNA*$_{ser}$ – *blpK* leader | JD0001 | UF_tRNAser | R_blpk_leader |
| | Spec fused-Dw HR of tRNA$_{ser}$ | JM1015 | F_spec | DR_tRNAser |
| | Mature sequence *psnE* | pUC57-PsnE | F_psnE (blpK leader) | R_pnsE (spec) |
| JD0014 | UP HR of *tRNA*$_{ser}$ – *blpK* leader | JD0001 | UF_tRNAser | R_blpk_leader |
| | Spec fused-Dw HR of tRNA$_{ser}$ | JM1015 | F_spec | DR_tRNAser |
| | Mature sequence *psnF* | pUC57-PsnF | F_psnF (blpK leader) | R_psnF (spec) |
| JD0015 | UP HR of *tRNA*$_{ser}$ – *blpK* leader | JD0001 | UF_tRNAser | R_blpk_leader |
| | Spec fused-Dw HR of tRNA$_{ser}$ | JM1015 | F_spec | DR_tRNAser |
| | Mature sequence *psnG* | pUC57-PsnG | F_psnG (blpK leader) | R_psnG (spec) |
| JD0016 | UP HR of *tRNA*$_{ser}$ – *blpK* leader | JD0001 | UF_tRNAser | R_blpk_leader |
| | Spec fused-Dw HR of tRNA$_{ser}$ | JM1015 | F_spec | DR_tRNAser |
| | Mature sequence *psnH* | pUC57-PsnH | F_psnH (blpK leader) | R_psnH (spec) |
| JD0017 | UP HR of *tRNA*$_{ser}$ – *blpK* leader | JD0001 | UF_tRNAser | R_blpk_leader |
| | Spec fused-Dw HR of tRNA$_{ser}$ | JM1015 | F_spec | DR_tRNAser |
| | Mature sequence *psnI* | pUC57-PsnI | F_psnI (blpK leader) | R_psnI (spec) |
| JD0018/JD0022 | UP HR of *tRNA*$_{ser}$ – *blpK* leader | JD0001 | UF_tRNAser | R_blpk_leader |
| | Spec fused-Dw HR of tRNA$_{ser}$ | JM1015 | F_spec | DR_tRNAser |
| | Mature sequence *psnJ* | pUC57-PsnJ | F_psnJ (blpK leader) | R_psnJ (spec) |
| JD0019/JD0023 | UP HR of *tRNA*$_{ser}$ – *blpK* leader | JD0001 | UF_tRNAser | R_blpk_leader |
| | Spec fused-Dw HR of tRNA$_{ser}$ | JM1015 | F_spec | DR_tRNAser |
| | Mature sequence *psnK* | pUC57-PsnK | F_psnK (blpK leader) | R_psnK (spec) |
| JD0020/JD0024 | UP HR of *tRNA*$_{ser}$ – *blpK* leader | JD0001 | UF_tRNAser | R_blpk_leader |
| | Spec fused-Dw HR of tRNA$_{ser}$ | JM1015 | F_spec | DR_tRNAser |
| | Mature sequence *psnL* | pUC57-PsnL | F_psnL (blpK leader) | R_psnL (spec) |
| JD0021/JD0025 | UP HR of *tRNA*$_{ser}$ – *blpK* leader | JD0001 | UF_tRNAser | R_blpk_leader |
| | Spec fused-Dw HR of tRNA$_{ser}$ | JM1015 | F_spec | DR_tRNAser |
| | Mature sequence *psnM* | pUC57-PsnM | F_psnM (blpK leader) | R_psnM (spec) |

#### 2. SUPPLEMENTAL FIGURES

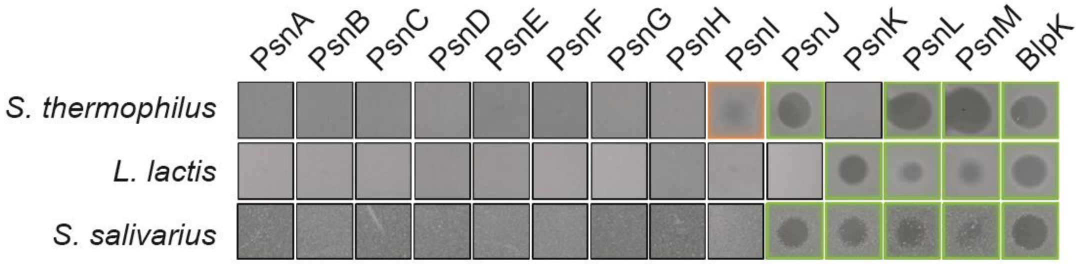

**Figure S1. Activity assays performed with mature bacteriocins produced *in vitro* by a cell-free system.** Spot-on-lawn assays showed that PsnJ, PsnK, PsnL, PsnM and BlpK are highly active (surrounded in green) and PsnI weakly active (surrounded in orange) against at least one of the three tested indicator strains (Table S5).

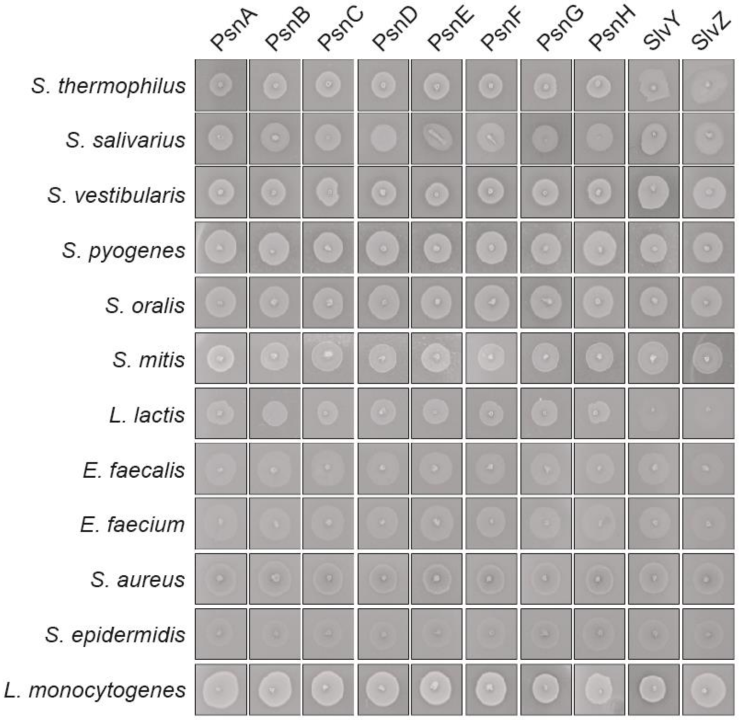

**Figure S2. Activity assays performed with single peptides of class IIb salivaricins produced *in vivo.*** The list of indicator strains is reported at Table S5.

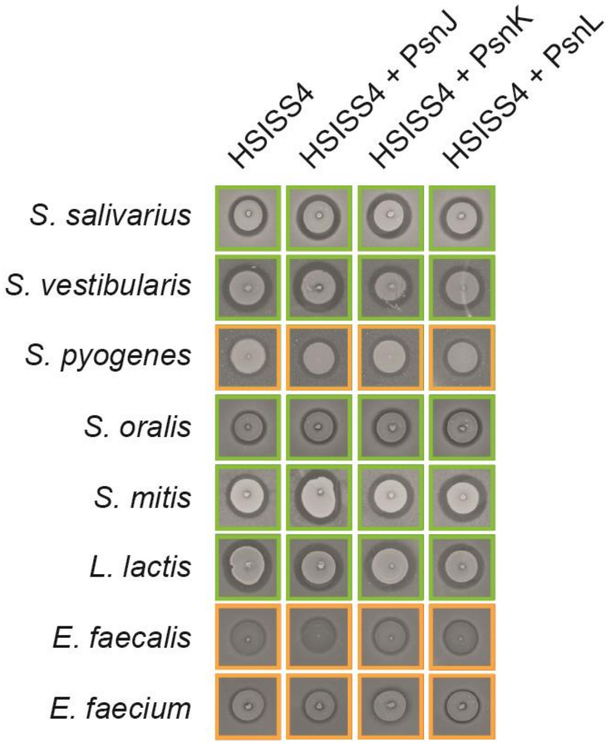

**Figure S3. Spot-on-lawn assays of HSISS4 derivatives producing PsnJ, PsnK, or PsnL against eight Gram-positive bacteria** (list of strains in Table S5). The HSISS4 wild-type strain is used as positive control. High and weak activities are surrounded by green and orange lines, respectively.
