## Supplemental Appendix S1 for "Uncovering the class II-bacteriocin predatiome in salivarius streptococci"

### Multiparametric bioinformatic script to identify class II bacteriocins.

This script was created by Julien Damoczi and Adrien Knoops.

The script is optimized for complete genomes. Nevertheless, it can be used with genomes fragmented in contigs. In this case, all headers (except the first one) need to be replaced by 4000 "A" to avoid false positives. Areas highlighted in yellow should be modified according to the analysis you wish to perform.

```
library(BiocManager)
library(msa)
library(Biostrings)
library(XML)
library(universalmotif)
library(writexl)
library(ggseqlogo)
library(systemPipeR)
library(seqinr)
library(writexl)
library(tidyr)
library(dplyr)
library(tidyverse)
library(Peptides)
library(alakazam)

workingDirectory <- "C:/Users/damoczi/OneDrive - UCL/Documents/Thèse/Résultat
s/Bioinfo/Script papier"

consensus <- readDNAStringSet(paste(workingDirectory,"consensus matrix ComR_B
ox salivarius.fasta", sep="/"))
msa_results <- msaClustalOmega(consensus, cluster="default", gapOpening="defa
ult", gapExtension="default")
PF_matrix <- consensusMatrix(msa_results)
PW_matrix <- PWM(PF_matrix)

#Create a list with all the genomes names

streptococcus <- readDNAStringSet(paste("C:/Users/damoczi/OneDrive - UCL/Docu
ments/Thèse/Résultats/Bioinfo/Script papier/S. salivarius2/complet/HSISS4.fna
"))

#search for motif with the PW matrix in the genome
```

```
plus_hits <- matchPWM(PW_matrix,streptococcus[[1]],min.score="75%")

df_hits_plus <- rbind(data.frame(start=start(plus_hits), end=end(plus_hits),
width=width(plus_hits),
                                 names="NZ_CP013216.1 Streptococcus salivariu
s strain HSISS4 chromosome, complete genome"))

results_sense <- data.frame("seqnames"=0,"subject_id"=0,"start"=0,
                            "end"=0,"width"=0,"strand"=0,"inframe2end"=0,"com
Rbox"=0,
                            "DNAseq"=0,"PROTseq"=0,stringsAsFactors=FALSE)

# Find the ORFs in the 4 kb after the comRbox
for (i in 1:nrow(df_hits_plus)){
  try(
    orf <- streptococcus[df_hits_plus$names[i]] %>%
      subseq(start = df_hits_plus$start[i], end = min(df_hits_plus$start[i]+4
000, streptococcus[df_hits_plus$names[i]][[1]] %>% length)) %>%
      chartr("N", "A",. ) %>%
      predORF(n=90, startcodon = c("ATG", "TTG", "GTG") , stopcodon = c("TAA"
, "TAG", "TGA"), type="df", mode="orf", strand="sense") %>%
      mutate(end = end+df_hits_plus$start[i]-1) %>%
      mutate(start = start + df_hits_plus$start[i]-1 ) %>%
      mutate(comRbox = df_hits_plus$start[i])
    , silent = T)


  if (orf$start %>% length() != 0){
    for (j in 1:max(min(90,orf$start %>% length()),1)){

      orf$DNAseq[j] <- streptococcus[df_hits_plus$names[i]][[1]] %>%
        subseq(start = orf$start[j],end = orf$end[j]) %>%
        as.character()

      orf$PROTseq[j] <- orf$DNAseq[j] %>%
        s2c %>%
        translate  %>%
        paste(sep="",collapse = "")
    }

    results_sense <- rbind(results_sense,orf)

  }
}

results_sense <- results_sense[-c(1),]
```

```r
Regulation_box <- data.frame("Regulation_Box")
for (m in 1:nrow(results_sense)) {
  start_pos <-  results_sense$comRbox[m]
  end_pos <- start_pos + 28
  upstream_seq <- subseq(streptococcus[1], end = end_pos , start = start_pos)
  Regulation_box[m] <- as.character(upstream_seq)
}
Regulation_box2 <- data.frame(t(Regulation_box))
results_sense <- cbind(results_sense, Regulation_box2)

#Reverse strand

streptococcus_rv <- reverseComplement(streptococcus)


#search for motif with the PW matrix in the genome
minus_hits <- matchPWM(PW_matrix, streptococcus_rv[[1]], min.score="75%")
df_hits_minus <- rbind(data.frame(start=start(minus_hits), end=end(minus_hits
), width=width(minus_hits),
                                  names="NZ_CP013216.1 Streptococcus salivari
us strain HSISS4 chromosome, complete genome"))

results_antisense <- data.frame("seqnames"=0,"subject_id"=0,"start"=0,
                                "end"=0,"width"=0,"strand"=0,"inframe2end"=0,
"comRbox"=0,
                                "DNAseq"=0,"PROTseq"=0,stringsAsFactors=FALSE
)

# Find the ORFs in the 4 kb after the ComRbox
for (i in 1:nrow(df_hits_minus)){
  try(
    orf <- streptococcus_rv[df_hits_minus$names[i]] %>%
      subseq(start = df_hits_minus$start[i], end = min(df_hits_minus$start[i]
+4000, streptococcus_rv[df_hits_minus$names[i]][[1]] %>% length)) %>%
      chartr("N", "A",. ) %>%
      predORF(n=90, startcodon = c("ATG", "TTG", "GTG") , stopcodon = c("TAA"
, "TAG", "TGA"), type="df", mode="orf", strand="sense") %>%
      mutate(end = end+df_hits_minus$start[i]-1) %>%
      mutate(start = start + df_hits_minus$start[i]-1 ) %>%
      mutate(comRbox = df_hits_minus$start[i])
    , silent = T)


  if (orf$start %>% length() != 0){
    for (j in 1:max(min(90,orf$start %>% length()),1)){

      orf$DNAseq[j] <- streptococcus_rv[df_hits_minus$names[i]][[1]] %>%
        subseq(start = orf$start[j],end = orf$end[j]) %>%
        as.character()
```

```
      orf$PROTseq[j] <- orf$DNAseq[j] %>%
        s2c %>%
        translate  %>%
        paste(sep="",collapse = "")
    }

    results_antisense <- rbind(results_antisense,orf)
  }
}
results_antisense <- results_antisense[-c(1),]

#add Regulation Box to the dataframe

Regulation_box <- data.frame("Regulation_Box")
for (m in 1:nrow(results_antisense)) {
  start_pos <-  results_antisense$comRbox[m]
  end_pos <- start_pos + 28
  upstream_seq <- subseq(streptococcus_rv[1], end = end_pos , start = start_p
os)
  Regulation_box[m] <- as.character(upstream_seq)
}
Regulation_box2 <- data.frame(t(Regulation_box))
results_antisense <- cbind(results_antisense, Regulation_box2)

#MERGING THE TWO RESULTS
results_final <- rbind(results_sense,results_antisense)


#GETTING ONLY THE ORF WITH THE MOTIF
gg_motifW <- subset(results_final,grepl("[L|V|M]....GG", results_final$PROTse
q))

#subselect sequence between 40 and 100 aa
gg_motifX <- subset(gg_motifW, width<300)
gg_motifY <- subset(gg_motifX, width>120)


#split leader and mature sequence
Want_name <- regmatches(gg_motifY$PROTseq[1], regexpr("GG", gg_motifY$PROTseq
[1]), invert = TRUE)

for (n in 1:nrow(gg_motifY)){
  Want_name[n] <- regmatches(gg_motifY$PROTseq[n], regexpr("GG", gg_motifY$PR
OTseq[n]), invert = TRUE)
}

Want_nameA <- unlist(Want_name)
Want_nameB  <- as.data.frame(matrix(Want_nameA, ncol = 2, byrow = TRUE))
```

```r
gg_motifY <-mutate(gg_motifY, Want_nameB)
GG <- "GG"
Want_nameD <- paste(gg_motifY$V1,GG, sep = '')
gg_motifY$V1 <- Want_nameD

#size leader sequence
for (n in 1:nrow(gg_motifY)){
  gg_motifY$Size_leader[n] <- width(gg_motifY$V1[n])
}

#keep only sequence with leader between 14 and 28 aa
gg_motifY <- subset(gg_motifY, Size_leader<29)
gg_motifY <- subset(gg_motifY, Size_leader>13)

#size mature sequence
for (n in 1:nrow(gg_motifY)){
  gg_motifY$Size_mature[n] <- width(gg_motifY$V2[n])
}

#keep only sequence with mature sequence between 25 and 70 aa
gg_motifY <- subset(gg_motifY, Size_mature<71)
gg_motifY <- subset(gg_motifY, Size_mature>24)

#GRAVY index
GRAVY <- data.frame("GRAVY")
for (n in 1:nrow(gg_motifY)) {
  GRAVY[n] <- gravy(gg_motifY$V2[n])
}

GRAVY2 <- data.frame(t(GRAVY))
gg_motifY <- cbind(gg_motifY, GRAVY2)

#Molecular weight
MW <- data.frame("MW")
for (n in 1:nrow(gg_motifY)) {
  MW[n] <- mw(gg_motifY$V2[n],monoisotopic = TRUE)
}

MW2 <- data.frame(t(MW))
gg_motifY <- cbind(gg_motifY, MW2)


#isoelectric_point
isoelectric_point <- data.frame("IP")
for (m in 1:nrow(gg_motifY)) {
  isoelectric_point[m] <- pI(gg_motifY$V2[m], pKscale= "EMBOSS")
}
isoelectric_point2 <- data.frame(t(isoelectric_point))
gg_motifY <- cbind(gg_motifY, isoelectric_point2)
```

```r
#Charge
Charge <- data.frame("Q")
for (m in 1:nrow(gg_motifY)) {
  Charge[m] <- charge(gg_motifY$V2[m], pH = 7)
}
Charge2 <- data.frame(t(Charge))
gg_motifY <- cbind(gg_motifY, Charge2)

colnames(gg_motifY) <- c('Seqnames','Subject_id','Start','End','Width','Stran
d','Inframe2end',
                         'ComRbox','DNAseq','PROTseq','Regulation box','LEADE
Rseq','MATUREseq','Size_leader',
                         'Size_mature','GRAVY','MW','Isoelectric_point','Char
ge')

new_order <- c('Seqnames','Start','End','Width','Regulation box','DNAseq','PR
OTseq','LEADERseq','MATUREseq','Size_leader',
                         'Size_mature','GRAVY','MW','Isoelectric_point','Char
ge')

gg_motifY <- gg_motifY[, new_order]


# EXPORTING RESULTS
write_xlsx(gg_motifY,paste(workingDirectory,"bacteriocins_of_HSISS4.xlsx", se
p="/"))
```
